# Islet macrophages shift to a reparative state following pancreatic beta-cell death and are a major source of islet IGF-1

**DOI:** 10.1101/480368

**Authors:** D. Nackiewicz, M. Dan, M. Speck, S. Z. Chow, Y.C. Chen, J. A. Pospisilik, C. B. Verchere, J. A. Ehses

**Author notes:** Corresponding Authors Dr. Jan A. Ehses, Twitter: @JanEhses, Dr. C. Bruce Verchere.

## Abstract

Macrophages play a dynamic role in tissue repair following injury. Here we found that following streptozotocin (STZ)-induced beta-cell death, mouse islet macrophages expressed increased *Igf1*, decreased proinflammatory cytokine expression, and transcriptome changes consistent with macrophages undergoing efferocytosis and having an enhanced state of metabolism. Macrophages were the major, if not sole, contributors to islet IGF-1 production. Adoptive transfer experiments showed that macrophages can maintain insulin secretion *in vivo* following beta-cell death with no effects on islet-cell turnover. IGF-1 neutralization during STZ-treatment decreased insulin secretion without affecting islet-cell apoptosis or proliferation. Interestingly, high fat diet (HFD) combined with STZ further skewed islet macrophages to a reparative state. Finally, islet macrophages from *db/db* mice also expressed decreased proinflammatory cytokines and increased *Igf1* mRNA. These data have important implications for islet biology and pathology and show that islet macrophages preserve their reparative state following beta-cell death even during HFD feeding and severe hyperglycemia.

## Introduction

Macrophages are versatile, plastic, innate immune cells essential to numerous biological processes. They participate in host defense, recognition of pathogens, initiation and resolution of inflammation, and maintenance of tissue homeostasis (Okabe and Medzhitov, 2016). Following tissue injury, macrophages are involved in the three main stages of tissue regeneration: inflammation, repair, and resolution. During the inflammatory phase, macrophages disrupt the basement membrane, secrete chemotactic factors to recruit inflammatory cells, and act as scavengers to phagocytose cellular debris. This is followed by a period of wound repair, where macrophages produce numerous growth factors, including insulin-like growth factor-1 (IGF-1), platelet-derived growth factors (PDGFs), and vascular endothelial growth factors (VEGFs) to stimulate blood vessel development and proliferation of neighboring parenchymal and stromal cells. Transforming growth factor β (TGF-β) is also produced during this stage and activates tissue fibroblasts to facilitate wound closure and extracellular matrix (ECM) deposition. In the last stage, macrophages assume an anti-inflammatory or pro-resolving phenotype characterized by anti-inflammatory cytokines (IL-10) and immune checkpoint inhibitor expression (PD-L2) (Vannella and Wynn, 2017).

In recent years, pancreatic islet macrophages have become increasingly well characterized in their resting state. Islet macrophages have an M1-like phenotype; they express *Il1b* and *Tnf* transcripts, MHC-II, present antigens to T cells, are negative for CD206/CD301, and are derived from definitive hematopoiesis (Calderon et al., 2015; Ferris et al., 2017). In the presence of aggregates of islet amyloid polypeptide (IAPP) (Masters et al., 2010; Westwell-Roper et al., 2016), or when exposed to toll-like receptor ligands (Nackiewicz et al., 2014), the proinflammatory state of islet macrophages is enhanced, leading to IL-1 secretion that causes beta-cell dysfunction (Nackiewicz et al., 2014; Westwell-Roper et al., 2014). In contrast, in transgenic models of pancreatic beta-cell regeneration, islet macrophages can produce factors that support beta-cell replication (Brissova et al., 2014; Riley et al., 2015).

Pancreatic beta-cell death is a feature of both type 1 and 2 diabetes, contributing to inadequate insulin secretion and clinical hyperglycemia in both diseases. In type 1 diabetes, apoptotic and necrotic beta-cell death occur. While the immunological consequences of apoptotic beta-cell death are unexplored, necrotic beta-cell death is thought to initiate or further enhance the activation of antigen-presenting cells in response to released beta-cell factors, causing T cell priming and activation, and promoting autoimmunity (Wilcox et al., 2016). In contrast, in type 2 diabetes apoptotic beta-cell death is mainly associated with disease pathology (Halban et al., 2014).

Very little is known about the dynamic role of islet macrophages following beta-cell death. We tested the hypothesis that islet macrophages could be skewed to a tissue-repair phenotype in response to beta-cell death, because apoptotic cells promote a tissue repair program in macrophages (Bosurgi et al., 2017) and other tissue macrophages have been shown to be locally programmed for silent clearance of apoptotic cells (Roberts et al., 2017). Here, we thoroughly characterized resident islet macrophage and recruited monocyte cell populations and gene signatures in response to STZ-induced cell death, in high fat diet (HFD)-STZ treated mice, and in *db/db* mice. Macrophages were the major source of IGF-1 protein within pancreatic islets and transcriptome changes post-STZ indicated an enhanced state of cellular metabolism and lysosome activity important in efferocytosis. Adoptive transfer of macrophages maintained circulating insulin levels following beta-cell death *in vivo*, and IGF-1 neutralization resulted in reduced 2^nd^ phase glucose-stimulated insulin secretion post-STZ *in vivo*.

## Results

### Islet macrophages in STZ-treated mice exhibit a gene set shift indicative of enhanced metabolism and lysosome activity and secrete IGF-1

Streptozotocin (STZ) is a toxin that specifically kills pancreatic beta-cells (Lenzen, 2008) and is commonly used to study islet inflammation. We recently used STZ to investigate the role of gp130 cytokine signaling in pancreatic alpha-cells in a model mimicking type 2 diabetes (STZ + high fat diet; (Chow et al., 2014)). When given at a repeated low dose (< 30 mg/kg) it induces mild effects on insulin secretion, glucose tolerance, and beta-cell mass (Chow et al., 2014). Here we used STZ to study the dynamic role of islet macrophages and monocytes following beta-cell death.

We initially analyzed islet macrophages and recruited monocytes at various time points post STZ treatment. Body weight and non-fasting blood glucose were unchanged up to 3 weeks post STZ (Suppl Fig. 1A-B). Islet CD45^+^ cells, islet macrophages, and recruited monocytes were increased following STZ, with significant increases starting at 1 week and peaking at 2 weeks (Fig. 1A-D). By 3 weeks, no further increases were detected (Fig. 1A-D). Changes in macrophage numbers correlated with gene expression changes, which showed the most difference at 2 weeks post STZ (Fig. 1E). At 2 weeks post STZ, *Tnf* mRNA expression was decreased and *Il1rn, Igf1,* and *Tgfbi* mRNA expression were increased in islet macrophages (Fig. 1E). No differences in mRNA expression of these genes were detected in recruited monocytes (Suppl Fig. 1C) and *Igf1* was consistently detected only in islet macrophages (see also Fig. 1E & Suppl Fig 1C).

**Figure 1:**
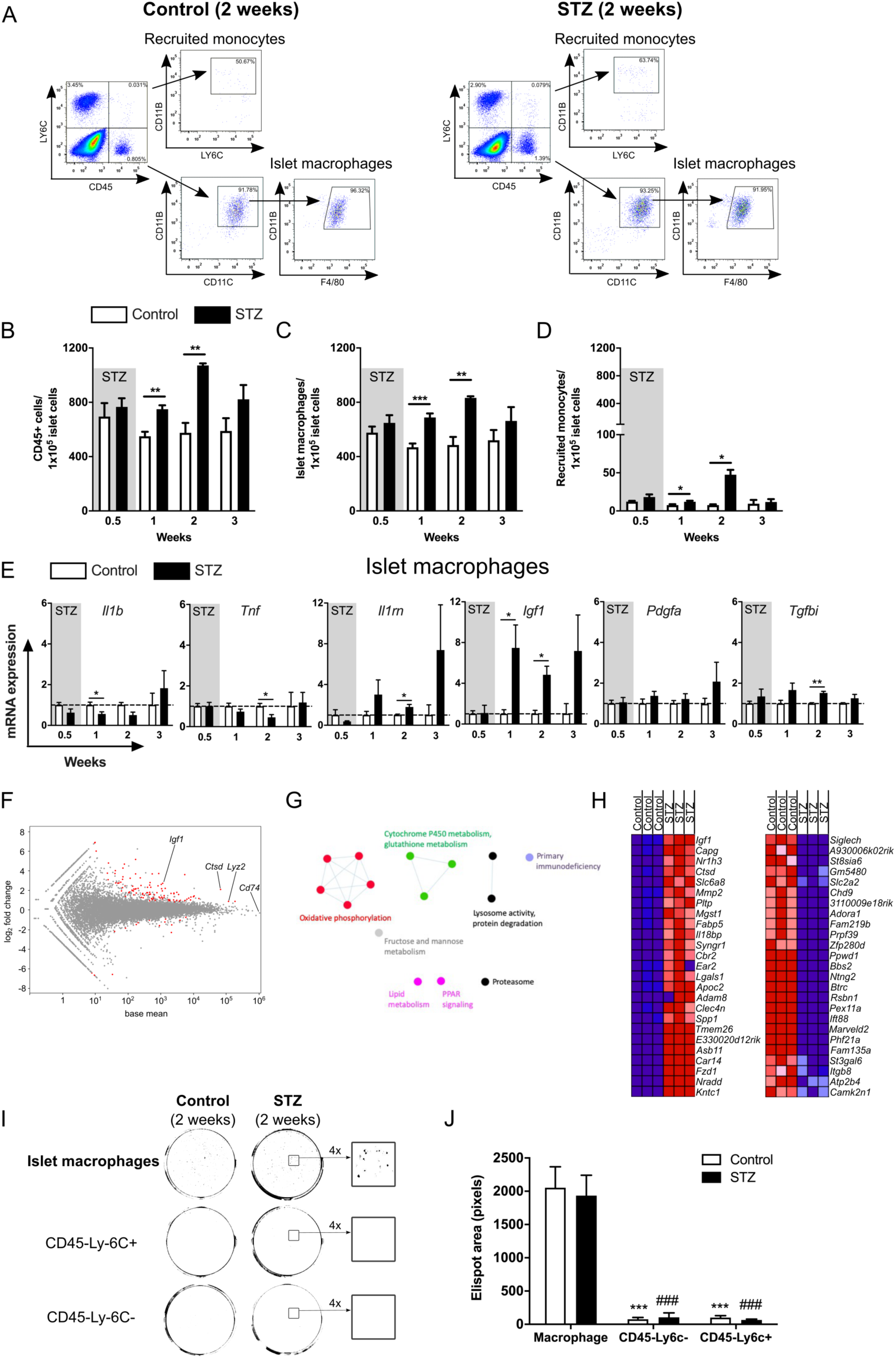
Islet macrophages in mice challenged with multiple low-dose STZ exhibit a gene shift towards enhanced metabolism and lysosome activity and secrete IGF-1. C57BL/6J male mice were given multiple low-dose STZ (30 mg/kg, 5 x daily i.p. injections) or acetate buffer as an injection control (referred to as “control”) at 16-20 weeks of age. (A) Representative flow cytometry plots and gating strategy for cell sorting of dispersed islets from mice treated with multiple low-dose STZ (right panel) or control treatments (left panel). Islets shown here were harvested 2 weeks after the first i.p. injection. Fractions of (B) CD45+ cells, (C) islet macrophages, (D) recruited monocytes. (E) qPCR of islet macrophages. Relative mRNA expression levels of *Il1b*, *Tnf*, *Il1rn*, *Igf1*, *Pdgfa*, and *Tgfbi* expressed as fold over islet macrophage control; n=3 for 0.5, 2, and 3-week treatments, and n= 5 for 1-week treatment. For each sorting sample (n), islets were pooled from 2-4 mice (average of 911 +/- 198 islets). (D-E) *p < 0.05, **p < 0.01, ***p < 0.001 STZ versus control, Student’s t test. (F-H) Transcriptome analysis of islet macrophages from mice treated with multiple low-dose STZ or control. Islets from 10 mice were pooled to create each sorting sample (n); n= 3, average of 2314 +/- 200 islets. (F) MA plot of islet macrophage gene expression post STZ with the mean of gene counts on the x-axis and log2 fold change of up- and down-regulated genes on the y-axis based on DEseq2 analysis. Significantly up- and down-regulated genes are shown in red (Log_2_ fold change > 1 and FDR < 0.05). (G) Enrichment map generated with Cytoscape of top-ranking clusters of genes enriched in STZ islet macrophages taken from GSEA analysis. Nodes represent gene-sets and edges represent mutual overlap. Highly redundant gene-sets are grouped together as clusters. Gene sets involved in similar biological processes are shown in the same color. (H) Heat map of GSEA results showing top 25 enriched genes in STZ (left panel) and top 25 enriched genes in control (right panel) islet macrophages (red, pink, light blue, dark blue corresponds to the range of expression values: high, moderate, low, lowest). (I) Representative IGF-1 ELISPOT images from cells sorted from dispersed islets. Two weeks after the first injection of buffer or multiple low-dose STZ, 2500 cells per group were sorted and analyzed by IGF-1 ELISPOT assay. On the right, outlined regions are enlarged 4 times. (J) Quantification of (I) IGF-1 ELISPOT area in pixels; n= 6, ***p < 0.001, islet macrophages from control mice versus non-immune cells; ###p < 0.001 islet macrophages from STZ treated mice versus non-immune cells, one-way ANOVA with Dunnett’s multiple comparisons test.

To obtain a broader unbiased view of the changes in islet macrophages following STZ, we performed transcriptome analysis of isolated islet macrophages at 2 weeks post STZ. The obtained gene expression profile was consistent with phagocytic immune cells, which are known to express high levels of *Cd74* (encoding part of MHC class II), *Lyz2* (also known as *LysM,* associated with lysozyme) and *Ctsd* (cathepsin D gene, associated with lysozyme; highlighted in Fig. 1F). 128 genes were found to be upregulated and 45 downregulated according to DEseq2 analysis (thresholds of Log_2_ fold change > 1 and FDR < 0.05). Gene Set Enrichment Analysis (GSEA) highlighted gene sets from various pathways enriched in macrophages from STZ-treated mice (Supplementary Table 1). There were no gene sets down-regulated with an FDR q value < 0.05. A Cytoscape enrichment map (Cline et al., 2007) of the main altered pathways highlighted multiple overlapping gene sets involved in increased oxidative phosphorylation, increased p450 metabolism, and increased lysosome and protein degradation activity (Fig. 1G). Increased lipid metabolism and PPAR signaling genes were also enriched in macrophages from STZ-treated mice. There were no inflammatory gene sets enriched and prototypical anti-inflammatory genes were not differentially expressed (*Il10, Arg1*, *Mrc1, Jag1, Il4ra*). A number of growth factors showed consistent up-regulation in all three samples based on FPKM values (*Igf1*, *Ngf*, *Pdgfc*, *Tgfbi*, *Vegfb*, *Vgf*). Interestingly, a heat map of the top 25 enriched genes resulted in *Igf1* having the highest score (3.47) among all enriched genes (Fig. 1H). Cathepsin D (*Ctsd*) and matrix metalloproteinase-2 (*Mmp2*), both involved in the proteolysis of IGF binding proteins (Mutgan et al., 2018), were also among the top 25 enriched genes. Taken together, the transcriptome of islet macrophages at 2 weeks post-beta-cell death indicates a heightened state of metabolism, increased lysosome activity, and a state characterized by strong induction of *Igf1*.

Isolated islet macrophages were confirmed to secrete IGF-1 protein *ex vivo*, while non-immune cells found in islets (CD45^-^Ly6C^+^ and CD45^-^Ly6C^-^ cells) did not (Fig. 1I-J). Finally, to confirm that STZ was causing beta-cell apoptosis, numbers of TUNEL^+^Insulin^+^ cells were assessed at 1 and 2 weeks post STZ. Streptozotocin induced a 3-fold increase in beta-cell apoptosis at 2 weeks (Suppl Fig. 1D-E).

### Islet macrophage depletion decreases *Igf1* expression following STZ-induced beta-cell death *ex vivo*

To isolate direct beta-cell STZ effects from indirect effects (e.g. elevated postprandial glucose) that may modulate islet macrophage gene expression *in vivo*, experiments were performed on isolated islets. Treatment of islets with increasing concentrations of STZ (0.25 - 4 mM) gave similar results to our *in vivo* studies, causing a decrease in *Il1b* mRNA expression, while significantly increasing *Tgfbi* and *Pdgfa* mRNA expression (Suppl Fig. 2A). Whole islet *Il1rn* and *Igf1* mRNA expression also tended to increase. Beta-cell *Ins1* and *Ins2* expression were unchanged, while *Pdx1* expression was increased by 4 mM STZ treatment (Suppl Fig. 2A). To determine the contribution of islet macrophages to these gene expression changes, islet macrophages were depleted using islets isolated from CD11c-DTR mice treated with diphtheria toxin (DT). CD11C+ cells are exclusively macrophages in the islet (Ferris et al., 2017). Macrophage depletion following DT was confirmed by flow cytometry (Suppl Fig. 2B). Depletion of islet macrophages completely abolished the STZ-induced increase in *Igf1* mRNA expression (Fig. 2A), while having no effect on beta-cell mRNA levels (*Ins1, Ins2, Pdx1*). Our data also indicate that macrophages are the main source of *Tgfbi* transcript levels in islets (Fig. 2A) which is interesting in the context of recent studies on TGFBI in islets (Han et al., 2011, 2014). As previously shown (Ferris et al., 2017; Nackiewicz et al., 2014; Westwell-Roper et al., 2014), islet macrophages are also the main contributors to islet *Il1b* and *Tnf* expression. STZ-treated islets had increased numbers of TUNEL^+^Insulin^+^ cells (Fig. 2B-C), with no effect on EdU^+^Insulin^+^ cells (Fig. 2D-E). Depletion of islet macrophages reduced EdU^+^Insulin^+^ cells (Fig. 2E).

**Figure 2:**
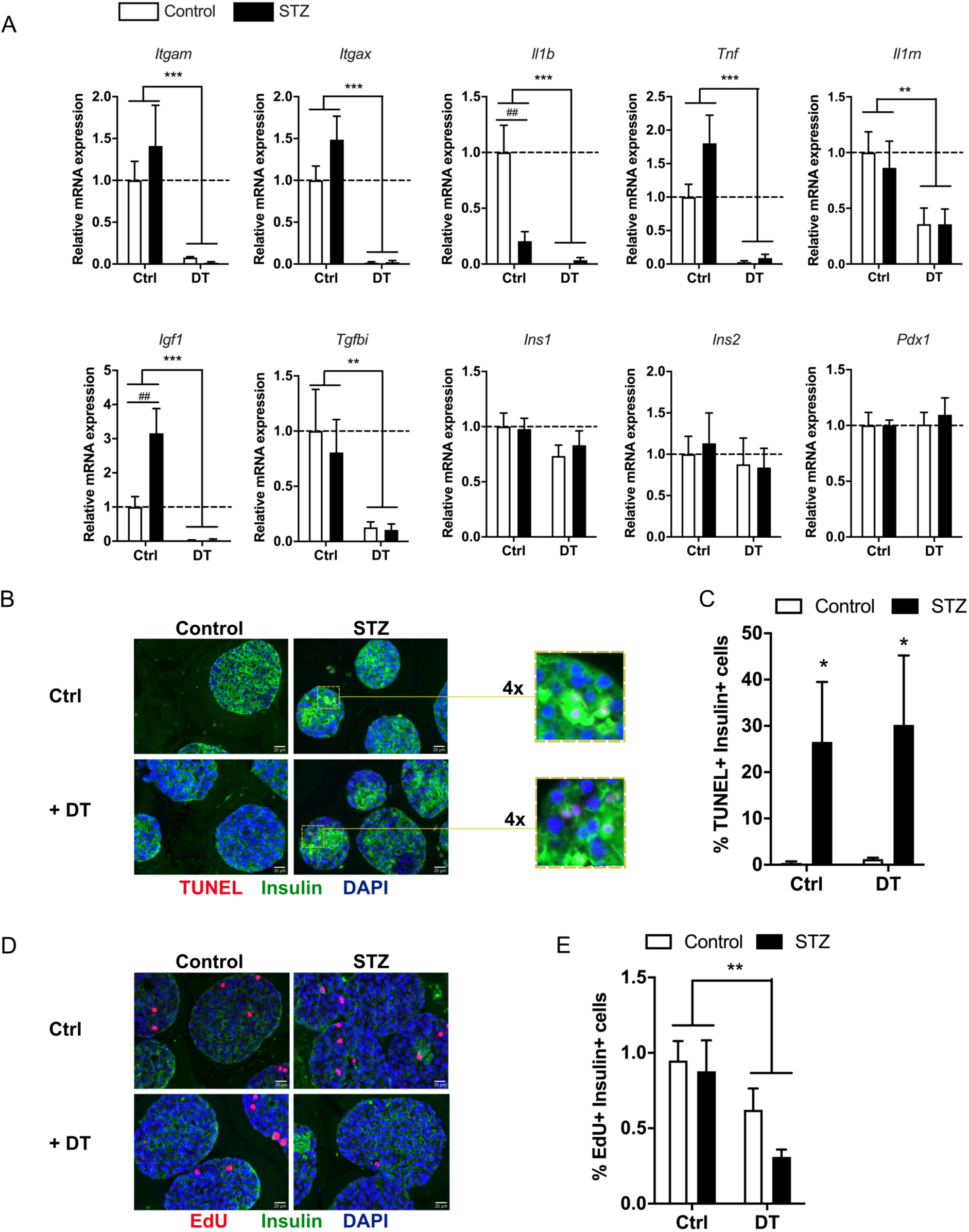
Islet macrophage depletion decreases *Igf1* expression following STZ-induced beta-cell death *ex vivo*. (A) qPCR of islets from male CD11c-DTR mice depleted (DT) or not (Ctrl) of islet macrophages followed by treatment with 4 mM STZ (STZ) or acetate buffer control (Control); n= 4-5, **p < 0.01, ***p < 0.001 DT versus Ctrl, ^##^p < 0.01 STZ versus Control, two-way ANOVA with Bonferroni’s multiple comparisons test. (B) Representative sections of TUNEL+ CD11c-DTR islets depleted (+ diphtheria toxin) or not (Ctrl) of islet macrophages followed by treatment with 4 mM STZ or acetate buffer control. Color scheme: DAPI-blue, insulin-green, TUNEL-red; colocalization of DAPI and TUNEL is shown in purple. Scale bar=20µm. (C) Quantification of (B). Between 544-4232 nuclei per section were counted. At least two sections from each sample (n) were counted; n=4, *p < 0.05 STZ versus control, two-way ANOVA with Bonferroni’s multiple comparisons test. (D) Representative sections of EdU-treated CD11c-DTR islets depleted (+ diphtheria toxin) or not (Ctrl) of islet macrophages followed by treatment with 4 mM STZ or acetate buffer control. Color scheme: DAPI-blue, insulin-green, EdU-red; colocalization of DAPI and EdU is shown in purple Scale bar=20µm. (E) Quantification of (D). Between 1412-3959 nuclei per section were counted. At least two sections from each sample (n) were quantified; n=3-5, **p < 0.01 DT versus Ctrl, two-way ANOVA with Bonferroni’s multiple comparisons test.

Finally, STZ did not increase *Igf1* mRNA expression or protein secretion, or affect proliferation or apoptosis in bone marrow derived macrophages (BMDMs; Suppl Fig. 3A-D). In summary, these data support the conclusion that macrophages are the major source of IGF-1 within islets and that beta-cell death directly stimulates islet macrophage *Igf1* mRNA expression.

### Macrophages and IGF-1 positively regulate insulin levels in mice following STZ

We next set out to further investigate the role of islet macrophages *in vivo*. Because islet macrophages already expressed elevated *Igf1* at 1 week post STZ, we depleted phagocytic cells with clodronate-loaded liposomes during, and immediately following, STZ (Fig. 3A). We recently used this protocol to deplete islet macrophages in mice (Nackiewicz et al., 2014; Westwell-Roper et al., 2014). There were no differences in body weight (Fig. 3B) between treatment groups, while non-fasting blood glucose was significantly elevated at the end of treatment in STZ-treated groups compared to their controls (Fig. 3C). Liposome-treated mice all tended to have decreased non-fasting insulin levels (Fig. 3D). Insulin secretion was significantly decreased in islets obtained from control mice treated with clodronate-loaded liposomes and in islets from STZ-treated mice (Fig. 3E), while islet insulin content was significantly reduced only in mice receiving STZ (Fig. 3F). Clodronate-liposome treated mice also tended to show worse glucose tolerance (Suppl Fig. 4A-B). No differences in TUNEL^+^ islet cells or pHH3^+^ islet cells were observed between groups (Suppl Fig. 4C-D). Effects of PBS-liposomes were consistent with a previously described effect of liposomes themselves on macrophage function (Ma et al., 2011; Pervin et al., 2016), or might have been due to macrophage depletion with PBS-liposomes (Weisser et al., 2011). These data show that islet macrophages help maintain beta-cell insulin secretion.

**Figure 3:**
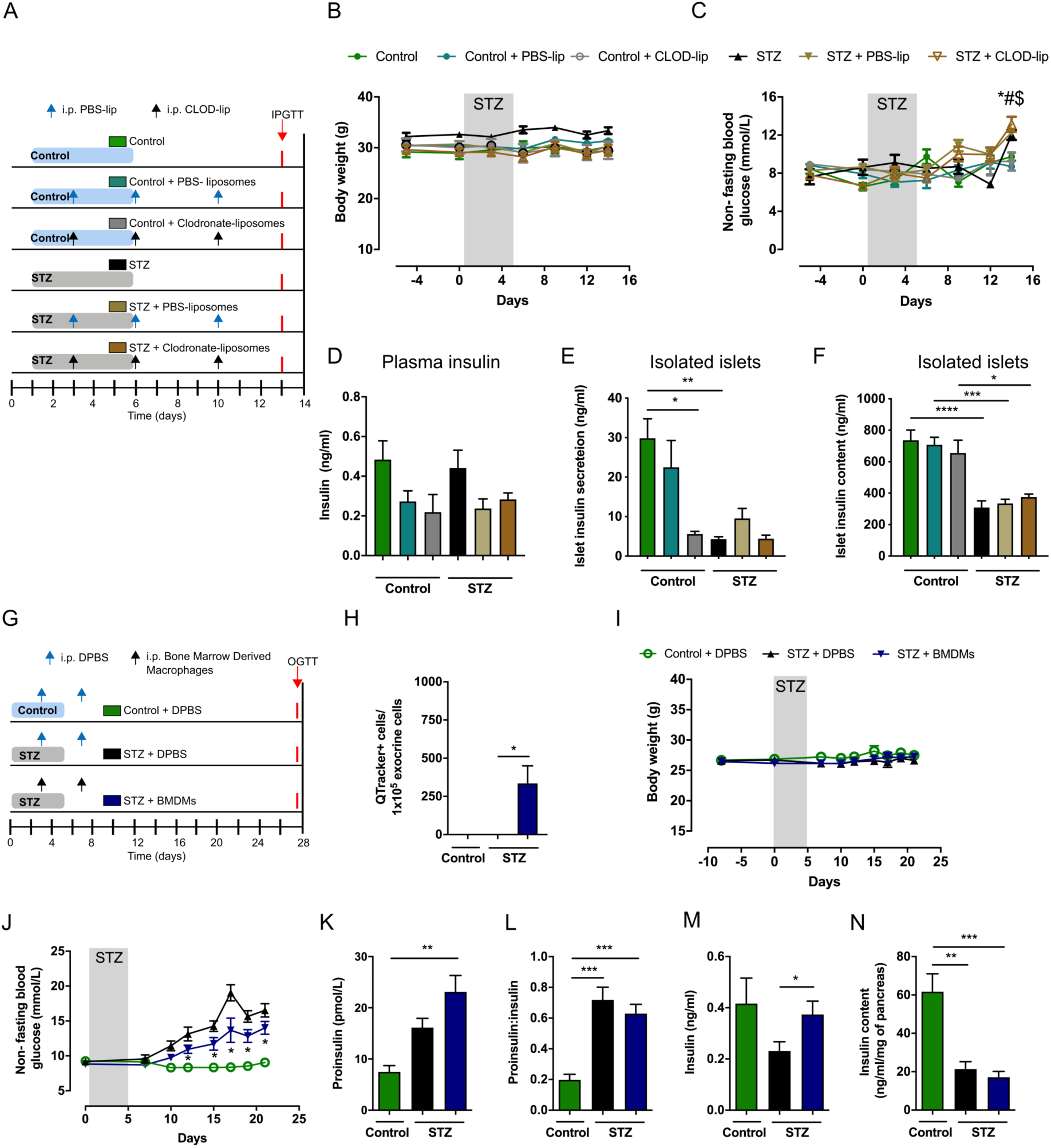
Macrophages positively regulate insulin levels in mice during multiple low-dose STZ. (A) Experimental design of macrophage depletion study displayed in (B-E). Multiple low-dose STZ (30 mg/kg, 5 x daily i.p. injections) or control (acetate buffer, 5 x daily i.p. injections) treatments were administered to C57BL/6J males two weeks before sacrifice. 200 µl of Clodronate-loaded liposomes (Clod-lip) or PBS-loaded liposomes (PBS-lip) were injected i.p. on day 3, 6, and 10 from the first dose of STZ/buffer. (B) Body weights; n=5 mice/control, control + PBS-lip, STZ + PBS-lip, STZ + CLOD-lip groups; n=4 mice/ STZ group, and n=3 mice/control + CLOD-lip group. (C) Non-fasting blood glucose measurements; n=5 mice/control, control + PBS-lip, STZ + PBS-lip, STZ + CLOD-lip groups; n= 4 mice/STZ group, and n=3 mice/control + CLOD-lip group, *p < 0.01 for control versus STZ, #p < 0.01 for PBS-lip versus STZ + PBS-lip, $ p < 0.01 for CLOD-lip versus STZ + CLOD-lip, two-way ANOVA with Tukey’s multiple comparisons test. (D) Cardiac puncture non-fasting insulin levels at the day of sacrifice; n=5 mice/control, control + PBS-lip, STZ + PBS-lip, STZ + CLOD-lip groups; n= 4 mice/STZ group, and n=3 mice/control + CLOD-lip group. (E) Islet insulin secretion in 2.8 mM glucose; n=5 mice/control, control + PBS-lip, STZ + PBS-lip, STZ + CLOD-lip groups; n= 4 mice/STZ group, and n=3 mice/control + CLOD-lip group. *p < 0.05, **p < 0.01, versus control, one-way ANOVA with Tukey’s multiple comparisons test. (F) Islet insulin content; n=5 mice/control, control + PBS-lip, STZ + PBS-lip, STZ + CLOD-lip groups; n= 4 mice/STZ group, and n=3 mice/control + CLOD-lip group. *p < 0.05, **p < 0.01, ***p < 0.001 versus control, one-way ANOVA with Tukey’s multiple comparisons test. (G) Design of experiments (H-N) involving adoptive transfer of bone-marrow derived macrophages (BMDMs). Multiple low-dose STZ (50 mg/kg, 5 x daily i.p. injections) or control (acetate buffer, 5 x daily i.p. injections) treatments were administered to C57BL/6J males four weeks before sacrifice. BMDMs that were starved of L929-conditioned medium were injected i.p. on day 3 and 7 from the first dose of STZ/buffer. (H) Qtracker positive cells in dispersed pancreas collected one day after second BMDM transfer; n= 4-5 mice/group. *p < 0.05, versus control Student’s t-test. (I) Body weights; n= 14-15 mice/group from 2 separate experiments. (J) Non-fasting blood glucose measurements; n= 14-15 mice/group from 2 separate experiments, *p < 0.01 STZ + BMDM versus STZ + DPBS, two-way ANOVA with Bonferroni’s multiple comparisons test. (K) Cardiac puncture serum non-fasting proinsulin levels at sacrifice; n= 6-12 mice/group. **p < 0.01, versus control one-way ANOVA with Dunnett’s multiple comparisons test. (L) Non-fasting proinsulin:insulin ratio at sacrifice; n= 6-12 mice/group. ***p < 0.001, versus control one-way ANOVA with Dunnett’s multiple comparisons test. (M) Cardiac puncture non-fasting serum insulin levels at sacrifice; n= 6-12, *p < 0.05, STZ + BMDM versus STZ + DPBS, Student’s t-test. (N) Pancreatic insulin content at sacrifice; n= 5, **p < 0.01, ***p < 0.001 versus control + DPBS, one-way ANOVA with Dunnett’s multiple comparisons test.

Next, we investigated if adoptively transferred macrophages could protect mice from STZ-induced hyperglycemia. Similar to islet macrophages, BMDMs are a rich source of IGF-1 protein (Suppl Fig. 3B) and secrete increased IGF-1 in response to phagocytosis of apoptotic cells (Han et al., 2016). We injected BMDMs intraperitoneally during, and immediately following, STZ (Fig. 3G). Injected macrophages homed to the pancreas (Fig. 3H), but could not be found in the spleen (data not shown). Macrophages had no effect on body weight (Fig. 3I), but significantly decreased non-fasting blood glucose (Fig. 3J). STZ-treated mice had elevated proinsulin levels and proinsulin:insulin ratios both with and without injected macrophages (Fig. 3K-L), but only mice receiving macrophages had significantly increased non-fasting insulin levels versus STZ controls (Fig. 3M). Insulin content was severely reduced due to STZ (Fig 3N), while no differences in TUNEL^+^ islet cells or EdU^+^ islet cells were observed between groups (Suppl Fig. 4E-F). Mice receiving macrophages also showed improved glucose tolerance (Suppl Fig. 4G). These data support the conclusion that macrophages that home to the pancreas can increase insulin secretion *in vivo* following STZ-induced beta-cell death and implicate a role for IGF-1.

Finally, because islet macrophages are the major, if not the sole source of IGF-1 in islets and its expression is upregulated following beta-cell death, we investigated if IGF-1 neutralization impacts glucose homeostasis during STZ-induced beta-cell death. We injected an IGF-1 antibody intraperitoneally during, and immediately following, STZ (Fig. 4A). IGF-1 neutralization had no effect on body weight, while non-fasting blood glucose was increased in the STZ + IgG group only (Fig. 4B-C). During a glucose challenge, only STZ + IGF-1 Ab mice had significantly impaired glucose tolerance versus IgG control mice (Fig. 4D-E). Significantly lower insulin levels at 30 minutes in STZ + IGF-1 Ab mice versus STZ controls coincided with the time point where the GTT curves separated (Fig. 4F,D). Similar to the adoptive transfer experiments, both groups of STZ-treated mice had elevated proinsulin levels and proinsulin:insulin ratios versus their controls (Fig. 4G-H), while non-fasting insulin levels tended to be lower in IGF-1 Ab treated mice (Fig. 4I). GH levels were unchanged between groups (Figure 4J). Finally, no differences in TUNEL^+^ islet cells or EdU^+^ cells were observed between groups (Suppl Fig. 5A-B). Thus, post STZ, IGF-1 signaling helps maintain 2^nd^ phase insulin secretion *in vivo*.

**Figure 4:**
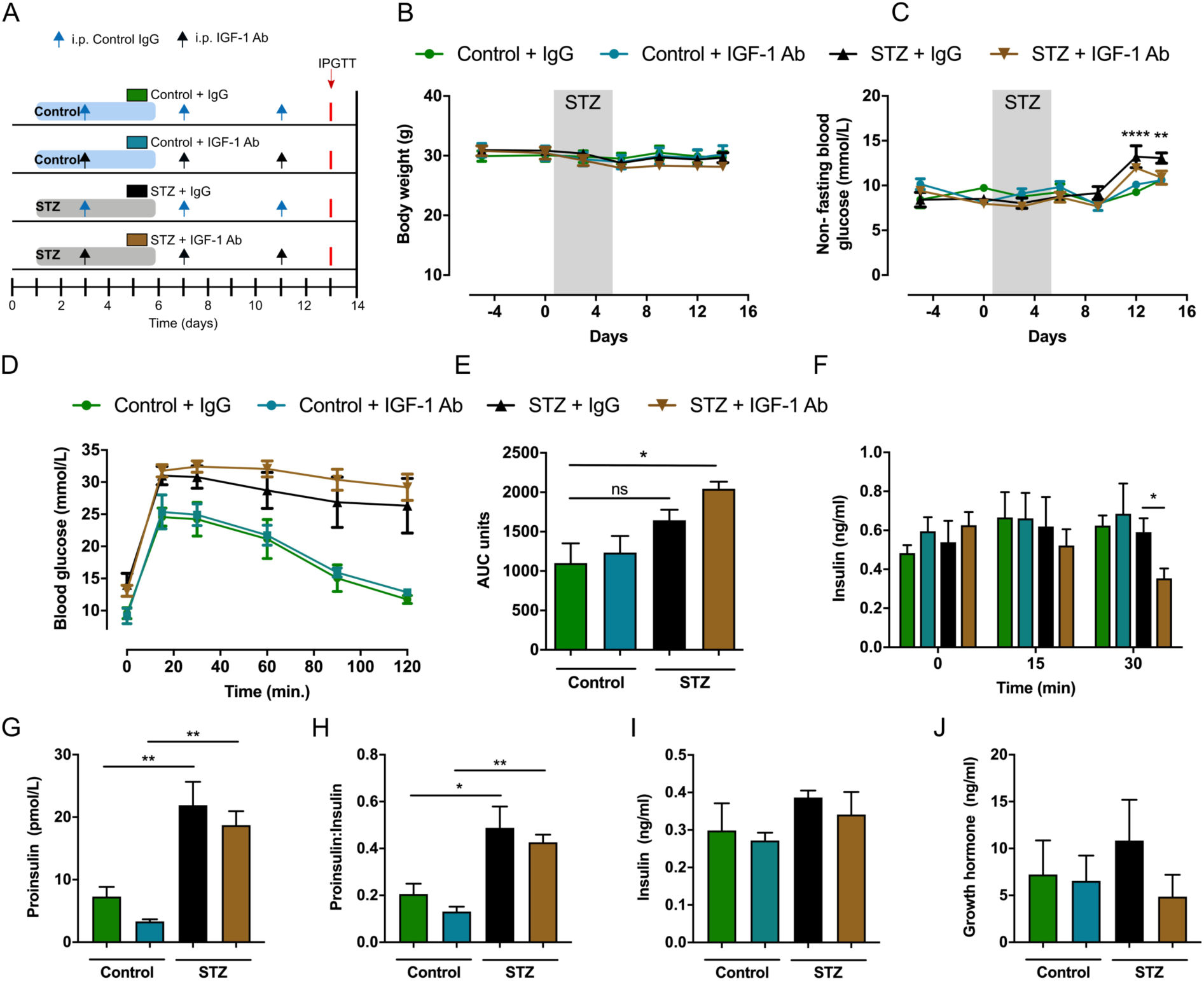
IGF-1 neutralization decreases insulin secretion in mice following multiple low-dose STZ. (A) Design of IGF-1 neutralization experiments (B-J). Multiple low-dose STZ (30 mg/kg, 5 x daily i.p. injections) or control (acetate buffer, 5 x daily i.p. injections) treatments were administered to C57BL/6J males two weeks before sacrifice. IGF-1 neutralizing antibody (0.1 µg/g body weight) or control IgG was injected i.p. on day 3, 7, and 11 following the first dose of STZ/buffer. (B) Body weights; n= 5 mice/group. (C) Non-fasting blood glucose measurements; n= 5 mice/group, ****p < 0.0001 STZ + IgG versus control + ctrl. IgG, two-way ANOVA with Dunnett’s multiple comparisons test. (D) Intraperitoneal glucose tolerance test (IPGTT, 1.5 g glucose/kg body weight) 13 days following the first dose of STZ or acetate buffer; n=5 mice, most of the blood glucose readings in STZ + IGF-1 neut. Ab group were above the range of the glucose meter and were recorded as 33.3 mmol/L- the highest reading within the range of the glucose meter. (E) Incremental area under the curve (AUC) for mice in (D); n=5, *p < 0.05 Control + IgG versus STZ + IgG, Kruskal-Wallis test with Dunn’s multiple comparisons test. (F) Serum insulin levels. Blood was collected at time 0, 15, and 30 min during IPGTT showed in (R); n=5, *p < 0.05 STZ + IgG versus STZ + IGF1 Ab at 30 min, Student’s t test. (G) Cardiac puncture non-fasting serum proinsulin levels from day 11-14; n= 5-7 mice/group from 2 separate experiments. **p < 0.01, STZ versus control one-way ANOVA with Tukey’s multiple comparisons test. (H) Non-fasting proinsulin:insulin ratio from day 11-14; n= 5-7 mice/group from 2 separate experiments. *p < 0.05, **p < 0.01 STZ versus control one-way ANOVA with Tukey’s multiple comparisons test. (I) Cardiac puncture non-fasting serum insulin levels from day 11-14; n= 8-9 from 2 separate experiments. (J) Cardiac puncture non-fasting serum growth hormone from day 11-14; n= 5.

### High fat diet further increases islet macrophage numbers and growth factor gene expression following beta-cell death

Beta-cell death and increased islet macrophages are usually associated with obesity in individuals with type 2 diabetes (Butler et al., 2003; Ehses et al., 2007). Therefore, we made mice obese by feeding them HFD. All mice were sacrificed at the same time point, 2 weeks post STZ, and at the same age (Fig. 5A). At 12 weeks, HFD and HFD+STZ mice had increased body weight compared to chow fed mice (Fig. 5B). Non-fasting blood glucose was increased in 12-week HFD+STZ mice (Fig. 5C), and glucose tolerance was impaired (Suppl Fig. 6A-B). Interestingly, numbers of CD45^+^ cells in islets were significantly increased in 12-week HFD+STZ mice versus STZ mice, due to increased numbers of islet macrophages and other CD45^+^Ly6C^+^CD11B^-^ cells (Fig. 5D-G and Suppl Fig. 6C-E). Numbers of CD45^-^Ly6C^-^ cells were also higher in 12-week HFD+STZ compared to STZ islets (Suppl Fig. 6D) while no differences in islet CD45^-^ Ly6C^+^ cell numbers were found (Suppl Fig. 6E). HFD did not increase islet monocytes versus STZ mice, similar to findings from Ying and colleagues (Ying et al., 2019).

**Figure 5:**
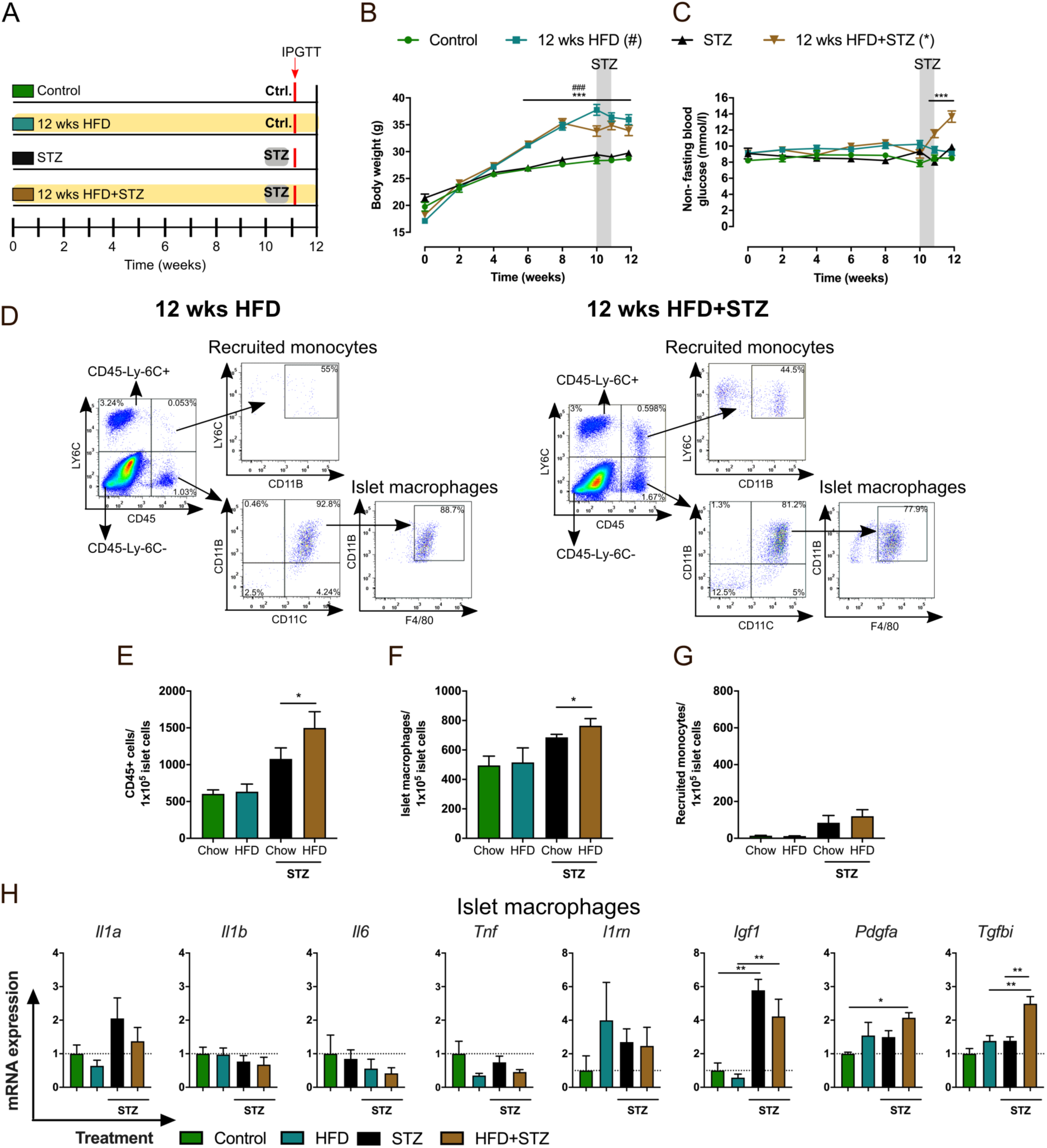
High fat diet further increases STZ-induced islet macrophages and growth factor gene expression. (A) Experimental design. C57BL/6J male mice were fed regular chow or HFD for 12 weeks. Multiple low doses of STZ (30 mg/kg, 5 x daily i.p. injections) or acetate buffer (referred to as “control”) were administered two weeks before sacrifice. (B) Body weights; n=12-22 mice/group, ^###/^***p < 0.001 HFD/HFD+STZ group versus control, two-way ANOVA with Dunnett’s multiple comparisons test. (C) Non-fasting blood glucose measurements; n=12-22 mice/ group, ***p < 0.001 HFD/HFD+STZ group versus control, two-way ANOVA with Dunnett’s multiple comparisons test. (D) Representative flow cytometry plots and gating strategy for cell sorting of dispersed islets from mice that received HFD for 12 weeks with acetate buffer injections (left panel) or with multiple low-dose STZ injections (right panel). Fractions of (E) CD45+ cells, (F) islet macrophages, (G) recruited monocytes from mice described in (A); n=5 for control, STZ groups; n=5-6 for 12 weeks HFD, 12 weeks HFD+STZ groups. *p < 0.05, Student’s t-test. (H) qPCR of islet macrophages. Relative mRNA expression levels of *Il1a*, *Il1b*, *Tnf*, *Il6*, *Il1rn*, *Igf1*, *Pdgfa*, and *Tgfbi* expressed as fold over islet macrophage control; n=4-6. For each sorting sample (n) islets from 3 mice were pooled together (average of 828 +/- 164 islets). *p < 0.05, **p < 0.01, test group versus indicated control, one-way ANOVA with Tukey’s multiple comparisons test.

Similar to islet macrophage gene expression post STZ alone, 12-week HFD+STZ macrophages also had increased *Igf1* mRNA expression versus HFD diet control (Fig. 5H). However, HFD did not further increase *Igf1* expression. Interestingly, HFD did further increase *Tgfbi* mRNA versus STZ alone (Fig. 5H). HFD+STZ macrophages also had significantly increased *Pdgfa* mRNA versus chow controls (Fig. 5H). No differences in mRNA expression of these genes were detected in recruited monocytes, CD45^-^Ly6C^-^ cells, or CD45^-^Ly6C^+^ cells (Suppl Fig. 6F-H). *Igf1* mRNA was consistently detected only in islet macrophages (see also Fig. 5H & Suppl Fig. 6F-H). In summary, HFD combined with STZ further increased numbers of islet macrophages and skewed islet macrophages to a state of increased growth factor expression.

### Islet macrophages in diabetic *db/db* mice express increased *Igf1* and decreased proinflammatory cytokines

Because STZ is a chemical toxin that might not be relevant for human disease, we also studied islet macrophages in a genetic rodent model of type 2 diabetes, the *db/db* mouse. At 6 weeks of age, *db/db* mice had elevated body weight, were hyperglycemic, hyperglucagonemic, and hyperinsulinemic compared to BKS controls (Fig. 6A-D). However, between 8-11 weeks of age insulin levels declined (Fig. 6D), indicative of beta-cell dysfunction and death (Medarova et al., 2005; Puff et al., 2011). Therefore, we investigated islet macrophages and monocytes at 8 and 11 wks of age. A trend towards increased numbers of CD45^+^ cells in *db/db* islets at 8 weeks of age was mainly due to significantly increased numbers of islet macrophages (Fig. 6E-G). Similar to islets post STZ, monocytes also tended to be increased (Fig. 6H). CD45^-^Ly6C^-^ cell numbers were increased and CD45^-^Ly6C^+^ cell numbers were significantly reduced (Suppl Fig. 7A-B).

**Figure 6:**
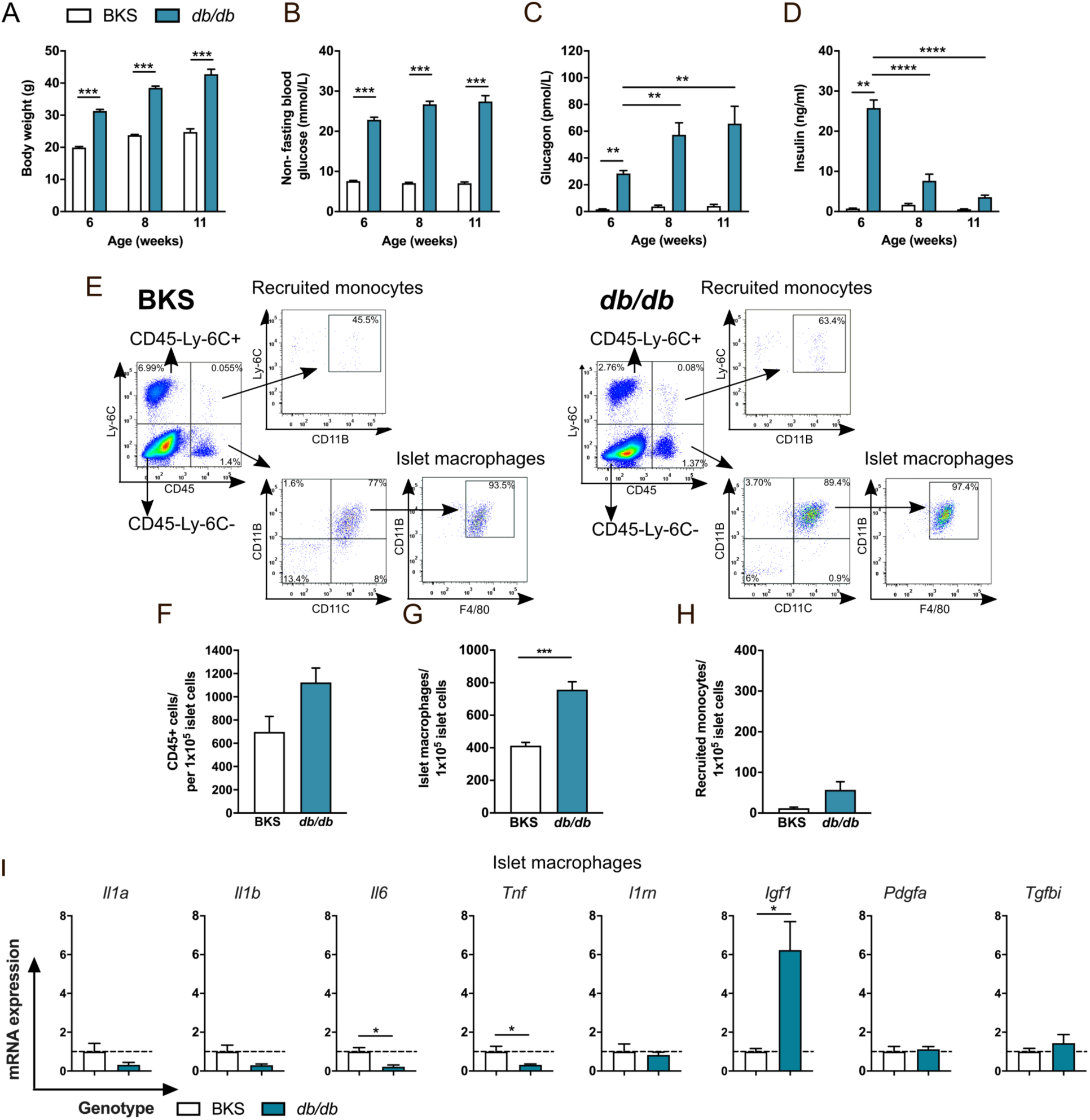
Islet macrophages in diabetic *db/db* mice express *Igf1* and decreased proinflammatory cytokines. (A) Body weight of 6-11-week-old male BKS and *db/db* mice. (B) Non-fasting blood glucose levels of 6-11-week-old BKS and *db/db* mice. (A-B) n=17-18 mice for 6-8-week-old groups, n=4 mice for 11-week-old group; ***p < 0.001 *db/db* versus BKS, Student’s t test. (C) Non-fasting glucagon levels; n=8 mice for 6-8-week-old groups, n=4 mice for 11-week-old group; **p < 0.01 6 wk old *db/db* versus BKS control, 8 and 11 wk old *db/db* versus 6 wk old *db/db*, one-way ANOVA with Tukey’s multiple comparisons test. (D) Non-fasting insulin levels; n=8 mice for 6-8-week-old group, n=3-4 mice for 11-week-old group; **p < 0.01, ****p < 0.0001 6 wk old *db/db* versus BKS control, 8 and 11 wk old *db/db* versus 6 wks old *db/db*, one-way ANOVA with Tukey’s multiple comparisons test. (E) Representative flow cytometry profiles and gating strategy for cell sorting of dispersed islets from 8-week-old BKS and *db/db* mice. Fractions of (F) CD45+ cells, (G) islet macrophages, (H) recruited monocytes in islets of 8-week-old BKS and *db/db* mice; (F-H) n=4, 2-4 mice pooled to obtain 556 +/- 52 islets per sample (n); ***p < 0.001 *db/db* versus BKS, Student’s t test. (I) qPCR of islet macrophages (G). Relative expression levels of *Il1a*, *Il1b*, *Tnf*, *Il6*, *Il1rn*, *Igf1*, *Pdgfa*, and *Tgfbi* expressed as fold control (BKS); n=4, 2-4 mice pooled per sample (n); *p < 0.05 *db/db* versus BKS, Student’s t test.

Assessment of cytokine (*Il1a, Il1b, Il6, Tnf, Il1rn*) and growth-factor (*Igf1, Pdgfa, Tgfbi*) mRNA expression in islet macrophages showed significantly reduced *Il6* and *Tnf* expression with 6-fold increased *Igf1* mRNA expression (Fig. 6I). No differences in mRNA expression of these genes were detected in monocytes (Suppl Fig. 7C). *Igf1* mRNA was consistently detected only in islet macrophages (see also Fig. 6I & Suppl Fig. 7C).

At 11 weeks of age, absolute numbers of CD45^+^ cells in islets also tended to be increased in *db/db* mice (Suppl Fig. 7D-E). Interestingly, this difference was no longer due to differences in islet macrophage or monocyte numbers (Suppl Fig. 7F-G), and was mainly due to an increase in other immune cell populations (CD45^+^LY6C^-^CD11B^-^ CD11C^-^, CD45^+^LY6C^+^CD11B^-^, Suppl Fig. 7D). Trends in other non-immune cells were similar to those seen at 8 weeks (Suppl Fig. 7H-I). Islet-macrophage cytokine and growth-factor mRNA expression showed a similar trend to data from 8-week-old *db/db* mice (Suppl Fig. 7J) with elevated *Igf1* mRNA.

In summary, similar to STZ-treated and HFD+STZ mice, islet macrophage numbers are increased in 8-week-old *db/db* mice, and gene expression indicates a state of increased *Igf1* expression, and decreased proinflammatory cytokine expression.

## Discussion

A number of studies, including our own, have shown that islets in humans with type 2 diabetes have increased numbers of macrophages (Ehses et al., 2007; Lundberg et al., 2017; Marchetti, 2016; Richardson et al., 2009). These cells express markers of both classical pro-inflammatory macrophages (CD68) and non-inflammatory macrophages (CD163) (Ehses et al., 2007). However, their functional role is unclear at present and can only be inferred from preclinical studies and clinical studies targeting the pro-inflammatory cytokine, IL-1 (Larsen et al, NEJM 2007; Everett et al., 2018).

Our findings here show that beta-cell death results in a dynamic increase in islet macrophage and recruited monocyte cells within 1 week, returning back to normal levels by 3 weeks. This is kinetically similar to effects seen in other tissues, such as cardiac tissue following injury (Walter et al., 2018). Selected pro-inflammatory, anti-inflammatory and growth factor genes showed changes at 2 weeks mainly indicative of a state of wound repair (Vannella and Wynn, 2017) and this was confirmed in our whole transcriptome analysis. Changes in macrophages paralleled significant increases in TUNEL+ beta-cells, potentially providing insight into how they shift to this state.

Phagocytosis of dead cells, also called efferocytosis, is known to induce an anti-inflammatory, reparative state in macrophages and has been increasingly studied in cardiovascular diseases (Brophy et al., 2017). Macrophages rapidly recognize and engulf apoptotic cells via so-called eat-me signals, the most fundamental of which is phosphatidylserine (PtdSer) (Lemke, 2019). Efferocytotic macrophages are characterized by a state of increased lysosome activity coupled with a state of increased energy needs (Henson, 2017; Voll et al., 1997). Recent studies have shown that fatty acid oxidation fuels the energy requirements of macrophages undergoing efferocytosis (Zhang et al., 2019). Macrophages undergoing phagocytosis of apoptotic cells are also known to secrete increased levels of IGF-1, which increases phagocytosis of local non-professional phagocytes and minimizes inflammation (Han et al., 2016). Our islet macrophage transcriptome data at 2 weeks post beta-cell death (increased expression of genes involved in oxidative phosphorylation, lysosome and protease activity, and lipid transport and oxidation, and increased *Igf1*) fits well with these known effects of efferocytosis on macrophages in other tissues. Future mechanistic studies should determine the phagocytic receptor responsible for shifting islet macrophages to this reparative state.

The energy requirements of macrophages undergoing efferocytosis may also help explain the changes seen in islet macrophages when beta-cell death was combined with HFD feeding. Enhanced skewing towards a reparative state under HFD feeding could be the result of an increased lipid energy source. Indeed, changes in cellular metabolism that lead to functional programming of phagocytic macrophages are a subject of considerable current interest; our data highlight potential pathways that could be targeted to promote islet macrophages with tissue-regenerative properties.

The liver is the main source of circulating IGF-1; however, it was recently proposed that macrophages could be the main source of extrahepatic IGF-1 (Gow et al., 2010). Indeed, numerous studies have identified macrophages as major producers of local tissue IGF-1 in the brain, in skeletal muscle, in the lung, and in various other tissues in studies investigating tumor associated macrophages in cancer biology (Forbes, 2016; Han et al., 2016; Ireland et al., 2016; Tonkin et al., 2015). Here we found that islet macrophages are the major source, if not the sole source of IGF-1 within pancreatic islets. This has important implications for islet biology.

IGF-1 has long been known to have a beneficial role in diabetes and cardiovascular disease (Higashi et al., 2019). Beta-cell IGF-1 overexpression protects from STZ-induced diabetes (George et al., 2002; Robertson et al., 2008) and exogenous IGF-1 protects NOD mice from developing type 1 diabetes likely via effects on T cells (Bergerot et al., 2008; Kaino et al., 1996). However, overexpression of IGF-1 locally within islets does not lead to an overt phenotype or effects on beta-cell mass suggesting its actions on beta-cells in diabetes are mainly indirect (George et al., 2002; Robertson et al., 2008). The only beta-cell effect observed in transgenic RIP-IGF-1 mice was increased 2^nd^ phase (30 minute) insulin secretion in response to a glucose challenge (Guo et al., 2005). This insulin secretory effect is in agreement with studies knocking out the IGF-1 receptor (IGF-1R) from beta-cells. The absence of a beta-cell IGF-1R did not impair beta-cell development or have effects on beta-cell mass, but it did result in increased fasting insulin levels and impaired 1^st^ and 2^nd^ phase insulin secretion in mice (Kulkarni et al., 2002; Xuan et al., 2002). These data agree well with our current findings, where the primary effect of IGF-1 on beta cells was to regulate 2^nd^ phase insulin secretion post beta-cell death. We propose that macrophages are the major local source of IGF-1 within pancreatic islets and that paracrine and autocrine effects of macrophage IGF-1 are critical in dampening islet inflammation and maintaining insulin secretion under pathological conditions (Fig. 7).

**Figure 7:**
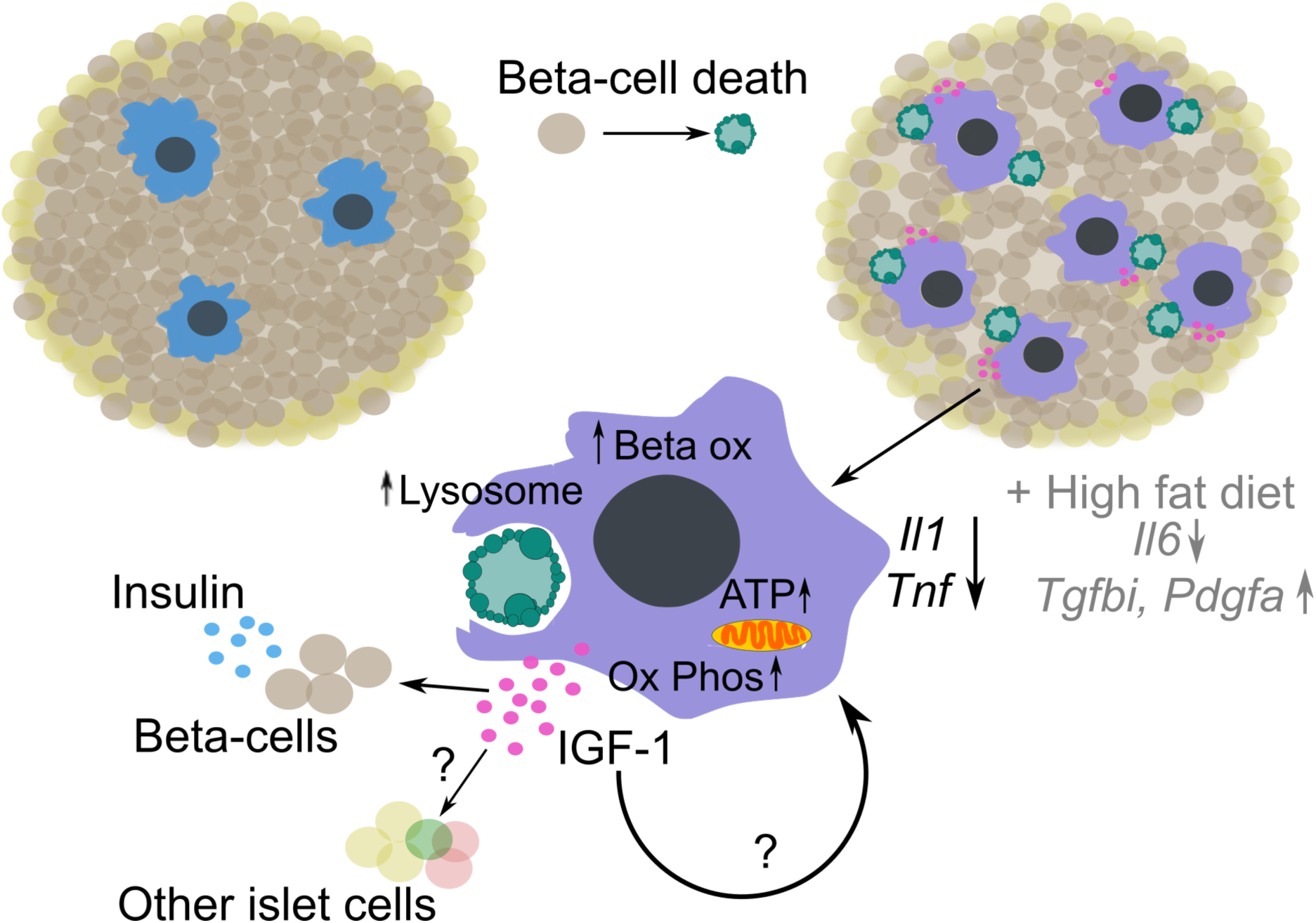
Schematic of the effects of beta-cell death on islet macrophages and IGF-1 secretion. Expression of genes involved oxidative phosphorylation, ATP production, lysosome βactivity, and lipid transport and oxidation are increased in islet macrophages following efferocytosis of dying beta-cells. During that process islet macrophages change their polarization state, decrease expression of pro-inflammatory cytokines, and increase IGF-1 secretion, even in the presence of HFD feeding or severe hyperglycemia. We propose that IGF-1 secreted by islet macrophages has paracrine and autocrine effects on beta-cells, other islet cell types, and macrophages themselves to limit inflammation and maintain beta-cell function under pathological conditions.

The present findings may seem counterintuitive given the established deleterious role of islet inflammation and IL-1 in humans with type 2 diabetes (Donath and Shoelson, 2011). Indeed, our own work has shown that islet amyloid polypeptide (IAPP) induces IL-1β in islet macrophages and toll-like receptor (TLR) ligands elevated in type 2 diabetes activate islet macrophages and have negative effects on beta-cell *Ins* gene expression (Nackiewicz et al., 2014; Westwell-Roper et al., 2014). However, in the absence of IAPP aggregates or certain TLR ligands islet macrophages clearly skew to a reparative phenotype following beta-cell death. We also observed a pro-resolution/anti-inflammatory phenotype in islet macrophages in *ex vivo* studies on GK rat islets isolated when beta cell function was declining *in vivo* (not shown).

When considering our data in the context of a chronic disease such as type 2 diabetes, it is important to keep in mind how dynamic inflammation and the function of a macrophage is. This is supported by our studies on the *db/db* mouse, where increased numbers of islet macrophages at 8 weeks where no longer evident at 11 weeks of age, when increases in other immune cell subsets were observed. It is also important to bear in mind that in our own work and that of others, global unbiased transcriptome analysis of islet macrophages was never conducted. It is entirely possible that islet macrophages exhibit a “mixed” pro-inflammatory and wound healing phenotype in type 2 diabetes in response to phagocytosis of dead beta-cells following cell death induced by IAPP and other factors (glucose, free fatty acids, ER stress, local and/or circulating cytokines). Regardless, our findings are in agreement with a recent study in *db/db* mice, where an NLRP3 inhibitor had no effects on glycemia (Kammoun et al., 2018). Ying and colleagues also found a mixed phenotype of islet macrophages during HFD feeding, not simply defined by M1 or M2 skewing (Ying et al., 2019). Similar to our findings, they also saw increased *Pdgfa* expression during HFD feeding (Ying et al., 2019).

A few limitations should be considered when interpreting our data. Currently available techniques do not allow manipulation of islet-specific macrophages *in vivo* due to lack of tissue-specific gene expression. Thus, macrophage depletion and adoptive transfer experiments should be interpreted with this in mind. To minimize this limitation, we made use of a toxin that specifically kills beta-cells (Lenzen, 2008), to enable study of the effects of macrophages in islet tissue. We also cannot completely exclude a role for liver IGF-1 in our neutralization experiments, despite effects on insulin secretion that were only evident following beta-cell death. GH levels were also not changed so there didn’t appear to be any impact of short-term IGF-1 neutralization on insulin sensitivity (Kim and Park, 2017).

Our data have important therapeutic implications relevant to type 2 diabetes. Despite the observance of a significant decrease in HbA1c, IL-1β inhibition has not shown prolonged effects on glycemic control in the studies conducted until now (Everett et al., 2018: Larsen et al, 2007). This could be due to the study population investigated until now or due to the dynamics of islet inflammation. Because inflammation is dynamic and islet macrophages can still change their phenotype to a reparative state even in the presence of HFD feeding or extreme hyperglycemia, increasing their beneficial effects might be an alternative approach to preserving functional beta-cell mass in type 2 diabetes.

## Methods

### Mice

BKS.Cg-Dock7m +/+ Leprdb/J (*db/db*), C57BLKS/J (BKS), B6.FVB-1700016L21Rik^Tg(Itgax-DTR/EGFP)57Lan^/J (CD11c-DTR) and C57BL/6J mice were purchased from Jackson Laboratory (Bar Harbor, ME). Mice were housed and bred (C57BL/6J and Cd11c-DTR) in the BC Children’s Hospital Research Institute Animal Care Facility in compliance with Canadian Council on Animal Care guidelines with a 12h light/12h dark cycle and fed *ad libitum* chow diet or where indicated high fat diet (58 kcal% fat w/sucrose Surwit Diet). Mature male mice were used in all experiments and were sacrificed between 14-20 weeks of age to ensure consistency between animal cohorts. BKS and *db/db* mice were 8-11 weeks old at sacrifice to coincide with declining beta cell function and death (Medarova et al., 2005; Puff et al., 2011). Mice were given either 30 mg/kg STZ or acetate buffer (control) i.p. for 5 consecutive days, or for 2 consecutive days if sacrificed on day 3. Following the first STZ or buffer injection mice were sacrificed as indicated in figures on day 3, 7 or 14 and islets were isolated by collagenase digestion. *In vivo* IGF-1 neutralization was achieved by injecting IGF-1 neutralizing antibody at 0.1 µg/g body weight i.p. Macrophages were depleted *in vivo* using clodronate loaded liposomes (or PBS loaded liposomes as controls) from Dr. Nico van Rooijen. Liposomes were allowed to reach room temperature, loaded into syringes, and the syringes inverted at least 10 times prior to injecting 200 µl of the solution i.p. per mouse. To adoptively transfer bone-marrow derived macrophages (BMDMs), mice were treated with either 50 mg/kg STZ or acetate buffer (control) i.p. for 5 consecutive days and ∼0.5 x 10^6^ BMDMs were injected i.p. on day 3 followed by injection of ∼1 x 10^6^ BMDMs i.p. on day 7 from the start of STZ/ buffer treatment. Mice were sacrificed on day 28 post STZ/buffer treatment. The numbers of animals studied are specified in each experiment. The University of British Columbia Animal Care Committee approved all animal studies.

### Mouse islet culture

After isolation, mouse islets were cultured in islet medium (RPMI 1640 medium (11.1 mM glucose, 2 mM L-Glutamine, Phenol Red) containing 10% FBS, 2 mM L-alanyl-L-glutamine dipeptide (GlutaMAX), 1% penicillin/streptomycin, 10 µg/ml of gentamicin) at 37°C in 5% CO_2_ and allowed to recover overnight prior to any *in vitro* experiments. For all *in vitro* experiments, 120-140 healthy-looking islets (round shape, absence of necrotic core, uniform brownish color) from multiple age matched males were pooled together for each n. To determine the optimal dose of STZ *in vitro*, islets were subjected to increasing concentrations of STZ in Krebs Ringer Bicarbonate Buffer (KRB) for 40 min and returned to mouse islet media for 48 h. Corresponding concentrations of acetate buffer were used as controls. To deplete CD11c cells from Cd11c-DTR islets, 10 ng/ml of diphtheria toxin was added to cultured islets for 24 h. Thereafter, islets were treated with 4 mM of STZ or acetate buffer in KRB for 40 min and returned to mouse islet media with 10 ng/ml of diphtheria toxin or vehicle (0.9% NaCl) for 48 h.

### Bone-marrow derived macrophages (BMDMs)

Bone marrow was spun down out of mouse femurs and tibias and BMDMs prepared as previously reported (Nackiewicz et al 2014). Red Blood Cell Lysis solution (0.155 M NH_4_Cl, 10 mM KHCO_3_, 0.127 mM EDTA) was used to deplete erythrocytes and the remaining cells were passed through 40 µm pore size strainers. Around 6 x 10^6^ cells were plated in 12 ml of DMEM supplemented with 1% penicillin/streptomycin, 10 mM HEPES, 10% FBS, and 15% L929-conditioned media in each 100 mm non-tissue culture treated dish and maintained at 37°C in 5% CO_2_. Fresh medium was added on days 3 and 5. On day 7, vigorous pipetting with Cell Dissociation Buffer was used to detach adherent BMDMs. Cells were plated in DMEM supplemented with 1% penicillin/streptomycin, 10% FBS at a density described in figures. After 24 h, BMDMs were used in experiments. BMDMs that were starved of L929 conditioned media (a source of macrophage colony-stimulating factor (M-CSF), nerve growth factor (NGF) and other undefined factors (Moore et al., 1980; Pantazis et al., 1977; Warren and Ralph, 1986)) for at least 24 h, were used for adoptive transfer experiments. Qtracker 655 Cell Labeling Kit (Invitrogen) was used to label BMDMs with fluorescent Qdot nanocrystals that allowed subsequent tracking of adoptively transferred cells in dispersed spleen, and exocrine pancreas by flow cytometry.

### Islet isolation

Mice anesthetized with isoflurane were sacrificed by cervical dislocation and islets isolated as previously reported (Nackiewicz et al 2014). After clamping the common bile duct, the pancreas was injected intraductally with approximately 2 mL of collagenase XI (1000 U/ml) in Hanks balanced salt-solution (HBSS) and placed in 50 mL tubes with an additional 3 mL of collagenase solution. The tube was incubated at 37°C for 14 minutes followed by gentle shaking to obtain a homogenously dispersed pancreas. Digestion was stopped with cold HBSS supplemented with 1 mM calcium chloride (CaCl_2_). Islets were washed two times in cold HBSS with CaCl_2_ and filtered through a 70 µM prewetted cell strainer. After flushing with 20 mL of HBSS with CaCl_2_, the strainer was turned upside-down over a Petri dish and rinsed with 10 mL of islet media to wash the islets into the dish. Islets were handpicked under the Nikon SMZ800 microscope into a fresh Petri dish with islet media.

### Physiological measurements

Non-fasting blood glucose levels were measured from tail bleeds at room temperature using a hand-held blood glucose meter and test strips (OneTouch® UltraMini®, OneTouch® Ultra®2, OneTouch® Ultra® Blue Test Strips, LifeScan Canada). Body weights were recorded at the same time. Mice were fasted 5 hours and injected intraperitoneally (i.p.) with 1.5 g glucose/kg of body weight or 1 g glucose/kg of body weight for i.p. glucose tolerance tests (IPGTT), or mice were given 2 g of glucose/kg of body weight for oral glucose tolerance tests (OGTT). Area under the curve (AUC) was calculated from baseline (time 0 min) for each animal and then used to determine the mean. Blood glucose levels during GTTs were measured from saphenous bleeds just before glucose injection and after 15, 30, 60 and 120 mins. Blood for serum insulin measurement was collected during 0, 15, and 30 min and measured using ELISA (Alpco). For plasma glucagon levels, aprotinin (250 kallikrein inhibitor units/mL plasma; Sigma-Aldrich) and dipeptidyl peptidase-4 inhibitor (50 μmol/L; Millipore) were added to the collection tubes and measured by ELISA (Mercodia). Insulin (Alpco), proinsulin (Mercodia), and growth hormone (Sigma-Aldrich) were measured by ELISA in sera from cardiac punctures.

### Immunocytochemistry

Isolated islets were fixed with 4% paraformaldehyde for 15 minutes at room temperature, washed with DPBS, set in agarose, embedded in paraffin and sectioned. Apoptosis was assessed by TUNEL staining with the In Situ Cell Death Detection Kit (Roche) according to the manufacturer’s directions. Proliferation was determined by EdU incorporation (islets were incubated with 10 µM EdU in islet media for 48 h or 72 h prior the fixation) and using the Click-iT™ EdU Alexa Fluor™ 594 Imaging Kit (Invitrogen) following the manufacturer’s directions or by anti-pHH3 antibody. Islet sections were blocked for 30 minutes at room temperature in 2% normal goat serum, incubated overnight at 4 °C with polyclonal guinea pig anti-insulin antibody (1:100 in 1% BSA in DPBS, DAKO) followed by 1 h room temperature incubation either with Alexa Fluor® 488 AffiniPure donkey anti-guinea pig or with DyLight™ 594 AffiniPure donkey anti-guinea pig secondary antibody (1:100 in 1% BSA in DPBS, Jackson ImmunoResearch Laboratories) and mounted using Vectashield with DAPI (Vector Laboratories). Imaging was acquired with a BX61 microscope and quantified using virtual slide microscope OlyVIA, ImageJ software and Image-Pro Analyzer.

### Pancreatic insulin content

Mice were sacrificed, and the pancreas was isolated. A small piece from the pancreatic tail was excised, weighed, homogenized in acid ethanol, and extracted overnight at 4°C. Samples were spun to remove debris. Supernatants were diluted, and insulin content measured by insulin ELISA (Alpco).

### Islet insulin secretion and content

Mice were sacrificed, and the islets were isolated by collagenase digestion as described above. After 24 h rest in islet media, 30 similarly-sized islets in duplicate per sample were pre-incubated for 30 min in a Krebs-Ringer Bicarbonate buffer containing 0.5% BSA, pH 7.2 (KRBH) and 2.8 mM glucose at 37°C. Subsequently, islets were transferred to fresh solution of 2.8 mM glucose in KRBH for 30 min at 37°C and supernatant was collected at the end of the incubation. Islet insulin content was extracted during overnight incubation in acid ethanol at 4°C. Insulin concentrations in supernatants were measured by insulin ELISA (Alpco).

### Flow cytometry and cell sorting

Islet macrophages were sorted as previously published (Nackiewicz et al., 2014) with additional antibodies used to differentiate recruited monocytes. Freshly isolated islets were dispersed in 0.02% Trypsin-EDTA for 3 minutes followed by up to 1 minute of pipetting under a stereomicroscope to obtain a single cell solution. Islet media was added to stop the reaction. Dispersed islets were washed with FACS buffer (1% heat inactivated FBS, 1 mM EDTA, 11 mM glucose in PBS). Cells were kept on ice and pre-incubated with Fc Block (1:100) for 5 minutes, followed by 30 min incubation with CD45-eFluor 450 (1:250; clone 30-F11), Ly-6C-APC (1:1,200; clone HK1.4), CD11b-PE (1:1,200; clone M1/700, F4/80-FITC (1:150; clone BM8), CD11c-PECy7 (1:150; clone N418), and the viability dye 7AAD (1:2,000). Unstained, single stains, and fluorescence minus one controls were used for setting gates and compensation, and cells were gated on single, live cells. The detailed gating strategy is shown in figures. A BD LSR II was used for flow cytometry and a BD Aria IIu instrument (BD Biosciences) was used for cell sorting with the help of the BC Children’s Hospital Research Institute FACS core facility.

### ELISPOT

Islets from 2 C57BL/6J males aged 16-20 weeks were pooled to obtain enough macrophages for one sample (n). Mice were treated 14 days earlier with 5 daily i.p. injections of 30 mg/kg STZ or acetate buffer and cell sorting was performed as described above. 2500 cells from each group were sorted, plated for 40 h on a 96-well PVDF plate pre-coated with IGF-1 capture antibody (Peprotech Inc.) and maintained in islet media at 37°C in 5% CO_2_. To detect secreted IGF-1, biotinylated anti-murine IGF-1 (Peprotech Inc.) and streptavidin-ALP were used according to the manufacturer’s instructions. BMDMs served as a positive control. The plate was developed with BCIP/NBT substrate and read on an ELISPOT reader AID Autoimmun Diagnostika GMBH (Germany). Spots were quantified with Image-Pro Analyzer. Pictures were converted to black and white and the number of pixels per well were measured. For visualization the black and white colors were inverted.

### Real-time PCR

Total RNA was isolated from whole islets and BMDMs using the NucleoSpin® RNA II kit (Macherey-Nagel), and from FACS-sorted cells using the RNeasy Micro Kit (Qiagen) following the manufacturer’s instructions. RNA was quantified using a NanoDrop 2000c (Thermo Scientific). cDNA from whole islets and BMDMs was generated using Superscript II (Invitrogen). cDNA from FACS-sorted cells was prepared using Superscript III (Invitrogen). Quantitative PCR was performed using PrimeTime primers and probes (Integrated DNA Technologies) and TaqMan MasterMix (ThermoFisher/ Applied Biosystems) in the ViiA7 Real-Time PCR System (ThermoFisher/ Applied Biosystems). Differential gene expression was determined by the 2^−ΔΔCt^ method with *Rplp0* used as a reference gene.

### Bulk RNA-seq

Male C57BL/6J mice aged 16-20 weeks were given either 30 mg/kg STZ or acetate buffer i.p. for 5 consecutive days. On day 14 following the first STZ/buffer injection mice were sacrificed and islets isolated. Islets from 10 mice were pooled per sample (n). Islets were hand-picked under the microscope, dispersed, and FACS-sorted as described above. Viable, single CD45+Ly6c-Cd11b+Cd11c+F4/80+ cells were sorted using a BD FACS Aria IIu directly into lysis buffer, and the RNeasy Plus Micro Kit from Qiagen was used to isolate total RNA. Total RNA quality control quantification was performed using an Agilent 2100 Bioanalyzer. All RNA samples had an RNA integrity number (RIN) ≥9.1. The NeoPrep Library Prep System (TruSeq Stranded mRNA Kit) from Ilumina was used for library preparation followed by sequencing using standard Illumina methods and Ilumina NextSeq500. RNA-Seq Alignment (BaseSpace Workflow) 1.0.0, TopHat (Aligner) 2.1.0, were used to map raw reads to the reference genome of Mus musculus (UCSC mm10). Cufflinks 2.2.1, BLAST 2.2.26+, DEseq2 (Love et al., 2014), VisR (Younesy et al., 2015), gene set enrichment analysis (GSEA 3.0, the pathway gene sets: gseaftp.broadinstitute.org://pub/gsea/gene_sets_final/c2.cp.v6.2.symbols.gmt (Mootha et al., 2003; Subramanian et al., 2005) were used to analyze the transcriptome. Cytoscape v3.7.0 with the enrichment map plugin was used to generate a gene set enrichment map based on GSEA analysis (Cline et al., 2007). A node cut-off Q-value of 0.05 and an edge cut-off of 0.5 were used.

### Statistical analysis

Data are reported as mean ± SEM. Statistical analysis with normality tests were performed, and graphs were created with GraphPad Prism version 7.00. Two-tailed Student’s t test was used when comparing two means. One-way ANOVA or Kruskal-Wallis test with Dunn’s multiple comparisons test was applied when comparing more than two groups, and two-way ANOVA was used when comparing two independent variables in at least two groups. To compare every mean with a control, Dunnett’s post-test was employed. Bonferroni’s or Tukey’s post-hoc test was used to compare different sets of means. Differences were considered significant at p<0.05. The n value and details on statistical analyses of each experiment are indicated in the figure legends.

### Data availability

All relevant data are available from the authors upon request. RNA-seq data have been deposited in the ArrayExpress database at EMBL-EBI (www.ebi.ac.uk/arrayexpress) under accession number E-MTAB-7234.

### Contact for reagent and resource sharing

Further information and requests for reagents may be directed to and will be fulfilled by the corresponding author, Dr. C. Bruce Verchere. Commercially available reagents are indicated in the Supplementary Table 2.

## Acknowledgments

We are grateful to Dr. Laura Sly, Dr. Francis Lynn, Dr. Heather Denroche, and Dr. Paul Orban from the BC Children’s Hospital Research Institute for helpful discussions and suggestions during the conduct of the study, to Mitsuhiro Komba from the Islet Core Facility, Dr. Lisa Xu from the Flow Core Facility, Dr. Jingsong Wang and Dr. Bao Ping Song from the Histology and Imaging Core Facilities, Dr. Derek Dai and Dr. Galina Soukhatcheva for their technical assistance, and to Ryan Vander Werff from the UBC Biomedical Research Centre Sequencing Core for help with RNA-seq sequencing. This work was supported by a grant from the Canadian Institutes of Health Research (CIHR). D.N. was supported by a CIHR-Vanier Canada Graduate Scholarship. C.B.V. is supported by an investigator award from BC Children’s Hospital and the Irving K. Barber Chair in Diabetes Research.

## Author contributions

Conceptualization, D.N. and J.A.E.; Methodology, D.N., M.D. and J.A.E; Investigation, D.N., M.D., M.S., S.Z.C., Y.C.C.; Formal analysis, D.N., J.A.P, J.A.E; Resources J.A.P., J.A.E., C.B.V.; Writing– Original Draft, D.N. and J.A.E.; Writing– Review & Editing, D.N., J.A.E., J.A.P., C.B.V.; Visualization, D.N., J.A.E.; Funding Acquisition, J.A.E. and C.B.V.; Supervision, J.A.E. and C.B.V.

## Supplementary figure legends

**Supplementary Fig. 1:**
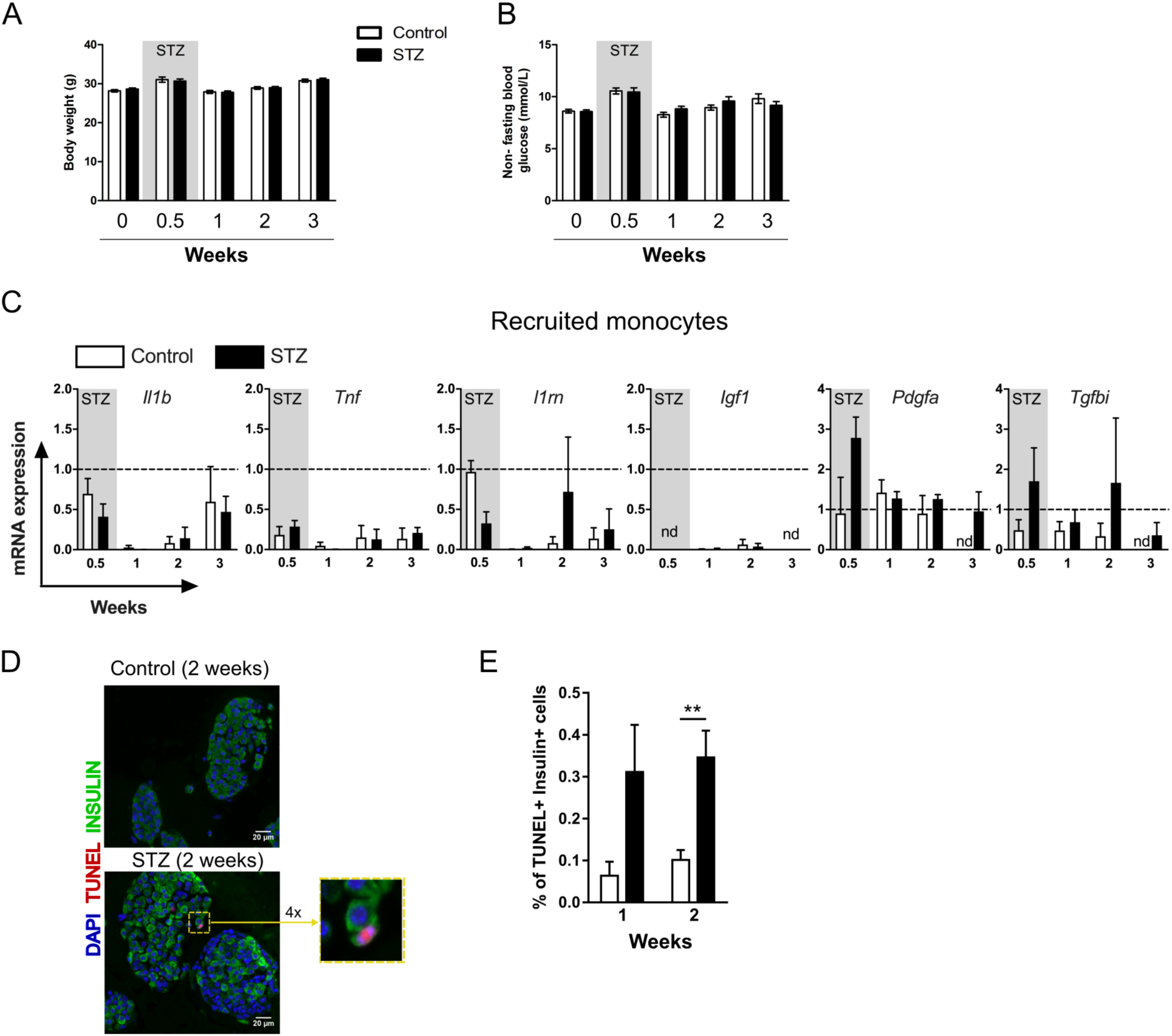
Body weight, glycemia, gene expression of recruited monocytes and beta cell apoptosis following STZ treatment *in vivo*. C57BL/6J male mice were given multiple low-dose STZ (30 mg/kg, 5 x daily i.p. injections) or acetate buffer as an injection control (referred to as “control”) at 16-20 weeks of age. (A) Body weights; n=9-11 mice/group. (B) Non-fasting blood glucose levels; n=9-11 mice/group. (C) qPCR of recruited monocytes 2 weeks post-STZ. Relative mRNA expression levels of *Il1β*, *Tnf*, *Il1rn*, *Igf1*, *Pdgfα*, and *Tgfβi* expressed as fold over islet macrophage control; n=3 for 0.5, 2, and 3-week treatments, and n= 5 for 1-week treatment. For each sorting sample (n), islets were pooled from 2-4 mice (average of 911 +/- 198 islets). (D) Representative sections of TUNEL+, insulin+ cells harvested from control or multiple low dose STZ treated mice 2 weeks post STZ. DAPI stain is shown in blue, insulin is green and TUNEL is visualized as a red color; scale bar=20µm. Colocalization of DAPI and TUNEL is shown in purple. On the right, outlined region is enlarged 4 times. (E) Quantification of TUNEL+, insulin+ cells harvested from control or multiple low-dose STZ treated mice after 1 and 2 weeks from the start of treatment. Between 567-9870 nuclei per section were counted; n=3-5, **p < 0.01 STZ versus control, Student’s t test.

**Supplementary Fig. 2:**
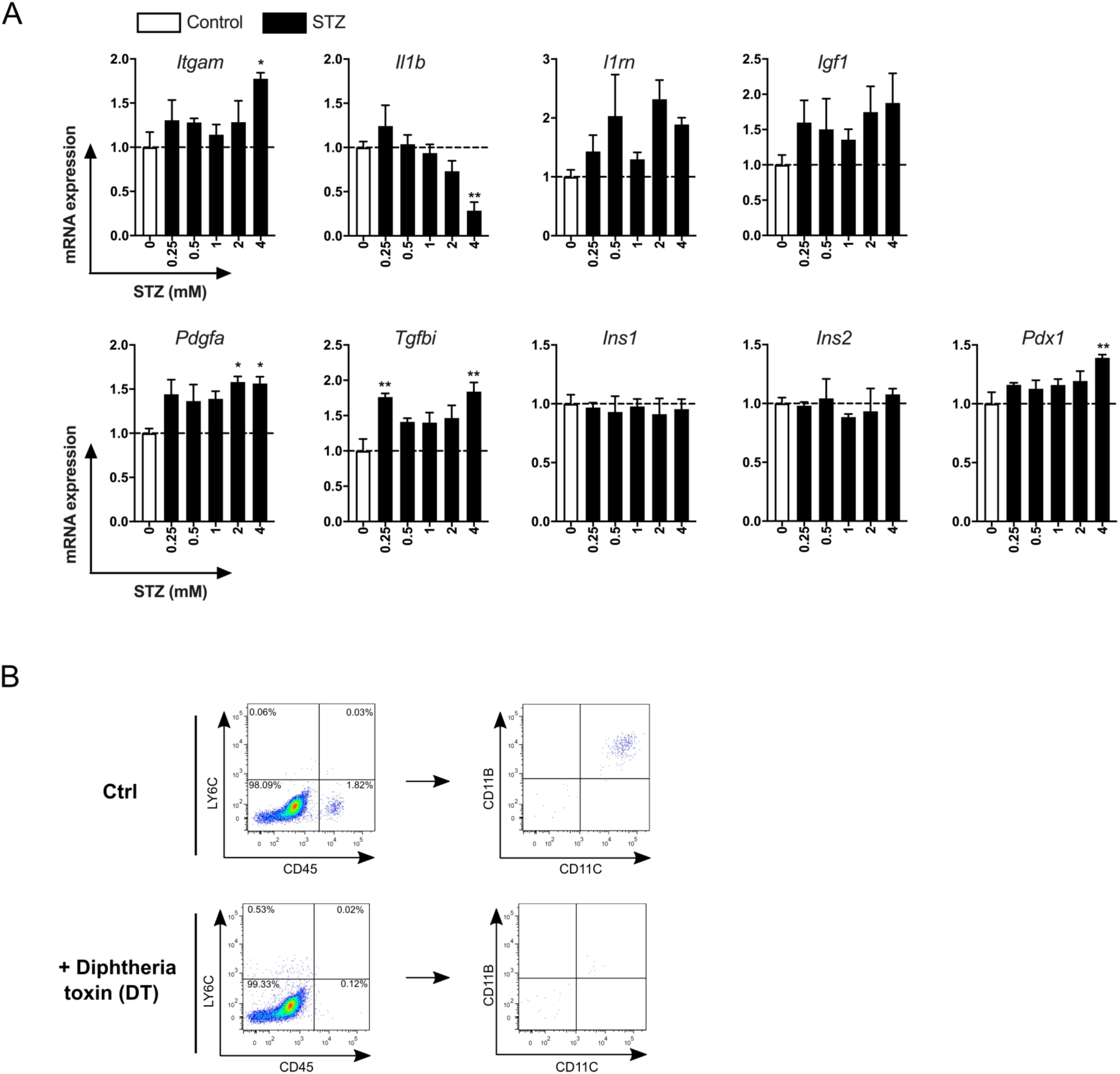
Dose response of STZ induced gene changes and flow cytometry of macrophage depletion in isolated islets. (A) qPCR of islets from male C57Bl/6 mice incubated *in vitro* with increasing doses of STZ for 40 min followed by 48 h recovery in islet media; n= 3, *p < 0.05, **p < 0.01 for STZ versus control, one-way ANOVA with Dunnett’s multiple comparisons test. (B) Representative flow cytometry plots of dispersed islets from male CD11c-DTR mice depleted or not of islet macrophages.

**Supplementary Fig. 3:**
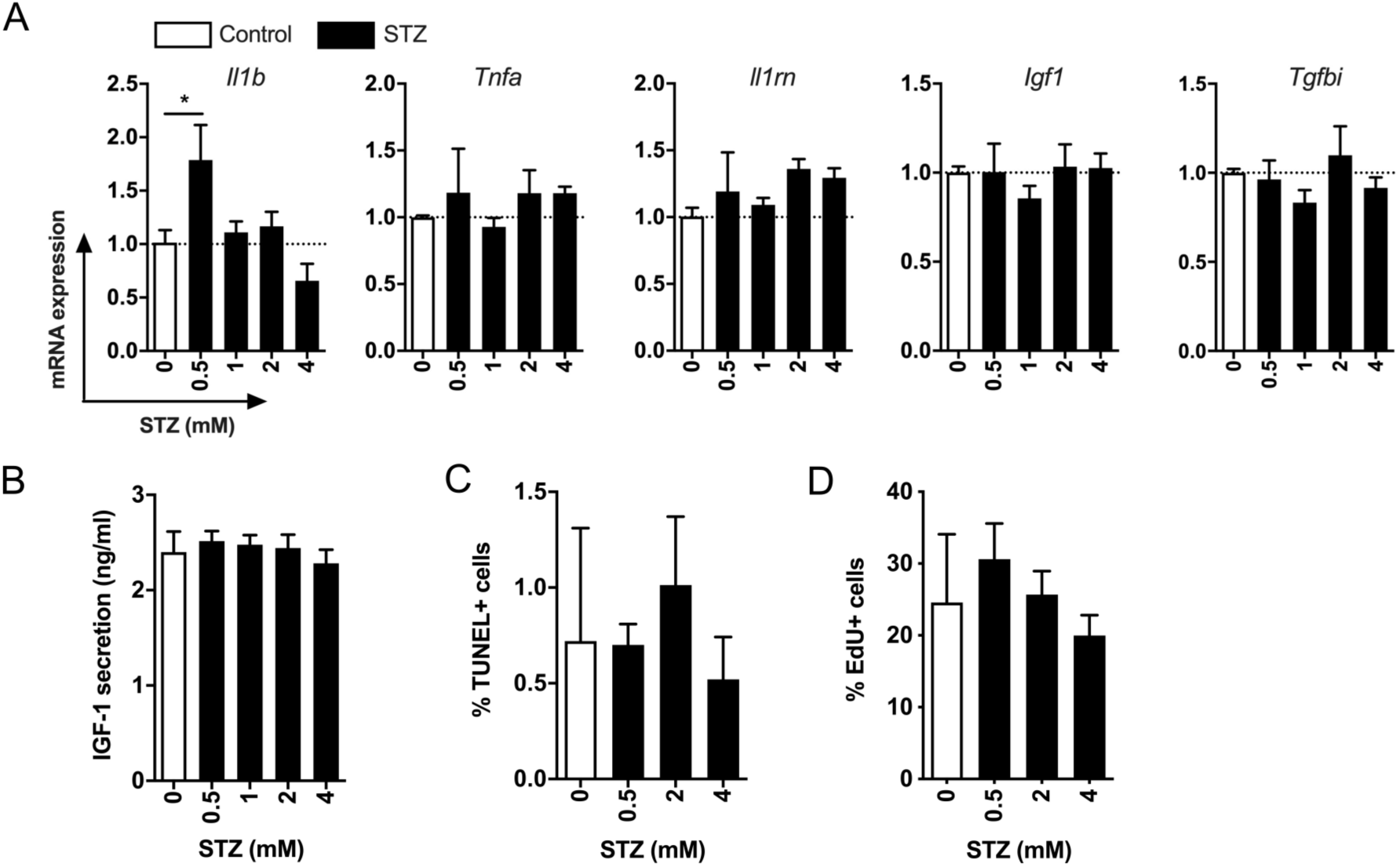
Bone marrow derived macrophages do not secrete IGF-1 following STZ treatment. (A) qPCR of BMDMs incubated *in vitro* with increasing doses of STZ for 40 min followed by 48 h recovery in islet media. Relative mRNA expression levels of *Il1β*, *Tnf*, *Il1rn*, *Igf1*, and *Tgfβi* shown as fold over control; n= 3, *p < 0.05 STZ versus 0 mM STZ, one-way ANOVA with Dunnett’s multiple comparisons test. (B) IGF-1 secretion from BMDMs incubated *in vitro* with increasing doses of STZ for 40 min followed by 48 h recovery in islet media; n= 3. (C) Quantification of TUNEL+ and (D) EdU+ BMDMs incubated *in vitro* with increasing doses of STZ for 40 min followed by 48 h recovery in islet media; n= 2.

**Supplementary Fig. 4:**
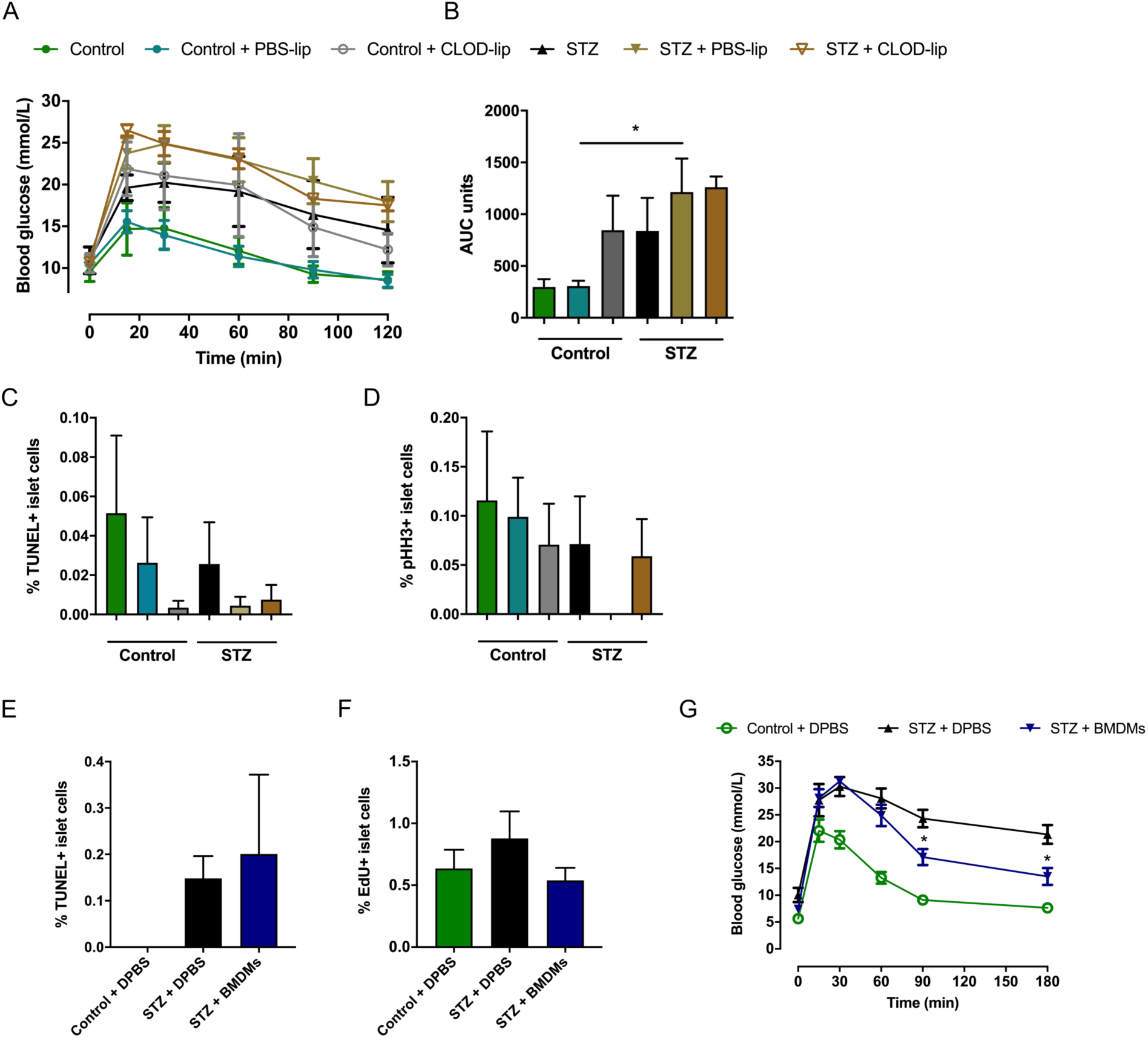
Glucose tolerance, islet-cell proliferation, and islet-cell apoptosis during macrophage depletion and adoptive transfer in the absence or presence of STZ. (A) Intraperitoneal glucose tolerance test (IPGTT, 1 g glucose/kg body weight) 13 days following the first dose of STZ/acetate buffer; n=5 mice/ control, control + PBS-lip, STZ + PBS-lip, STZ + CLOD-lip groups; n=4 mice/ STZ group, and n=3 mice/control + CLOD-lip group. (B) Incremental area under the curve (AUC) for mice in (A); n=5 mice/control, control + PBS-lip, STZ + PBS-lip, STZ + CLOD-lip groups; n= 4 mice/STZ group, and n=3 mice/control + CLOD-lip group, *p < 0.05 STZ versus control, one-way ANOVA with Tukey’s multiple comparisons test. (C) Quantification of TUNEL+ islet cells and (D) pHH3+ islet cells harvested from control or multiple low-dose STZ treated mice 2 weeks from the start of treatment. Between 394-16144 nuclei per section were counted; n=3-5, One-way ANOVA with Tukey’s multiple comparisons test. (E) Quantification of TUNEL+ islet cells and (F) EdU+ islet cells in pancreatic sections from control or multiple low-dose STZ (50 mg/kg) treated mice 28 days from the start of the STZ/control treatment. EdU (1 mg) was injected daily i.p. for the last five days before the sacrifice. Between 370-2615 islet cells per section were counted; n=3-5, One-way ANOVA with Tukey’s multiple comparisons test. (G) Oral glucose tolerance test (OGTT, 2 g glucose/kg body weight) 25-26 days following administration of the first dose of STZ or acetate buffer; n= 5-6 mice, *p < 0.05, STZ + BMDM versus STZ + DPBS, two-way ANOVA with Bonferroni’s multiple comparisons test.

**Supplementary Fig. 5:**
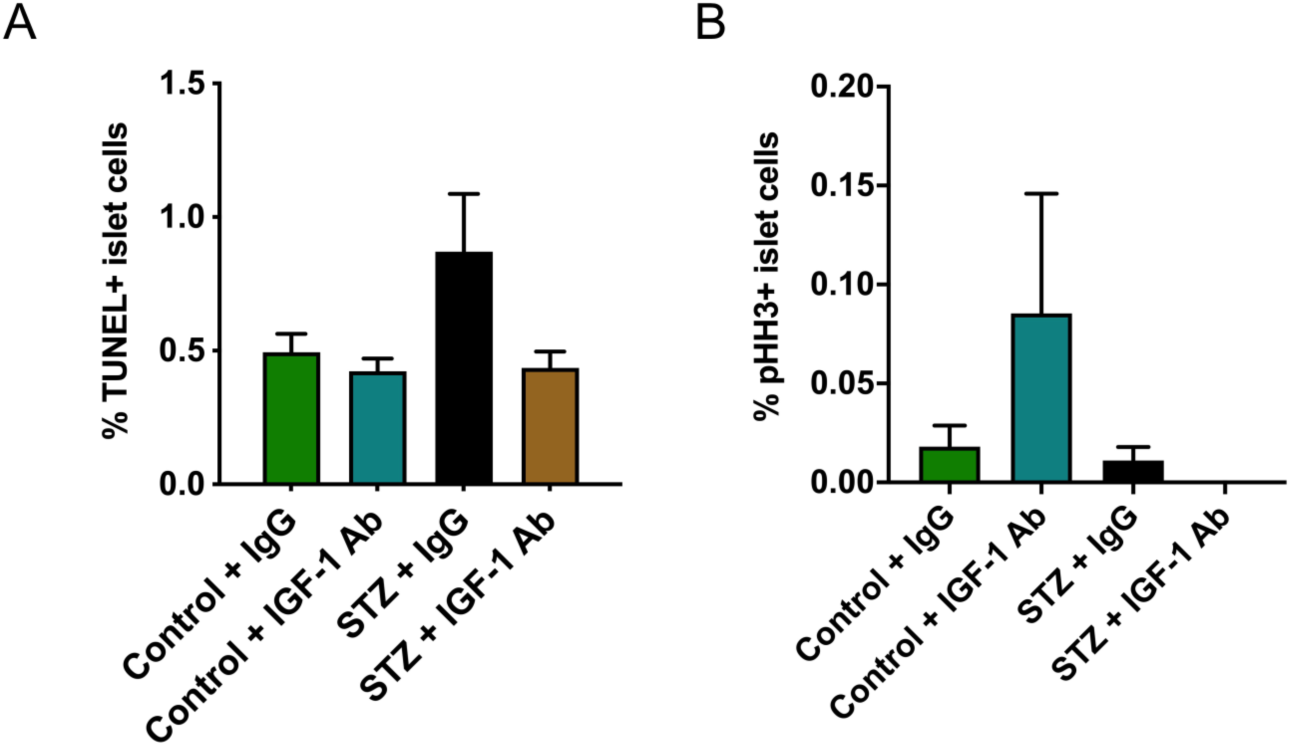
Islet-cell apoptosis and proliferation during IGF-1 neutralization in the absence or presence of STZ. (A) Quantification of TUNEL+ islet cells and (B) pHH3+ islet cells harvested from control or multiple low-dose STZ treated mice 2 weeks from the start of treatment. Between 931-13095 nuclei per section were counted; n=3-5, **p < 0.01 STZ versus control. One-way ANOVA with Tukey’s multiple comparisons test.

**Supplementary Fig. 6:**
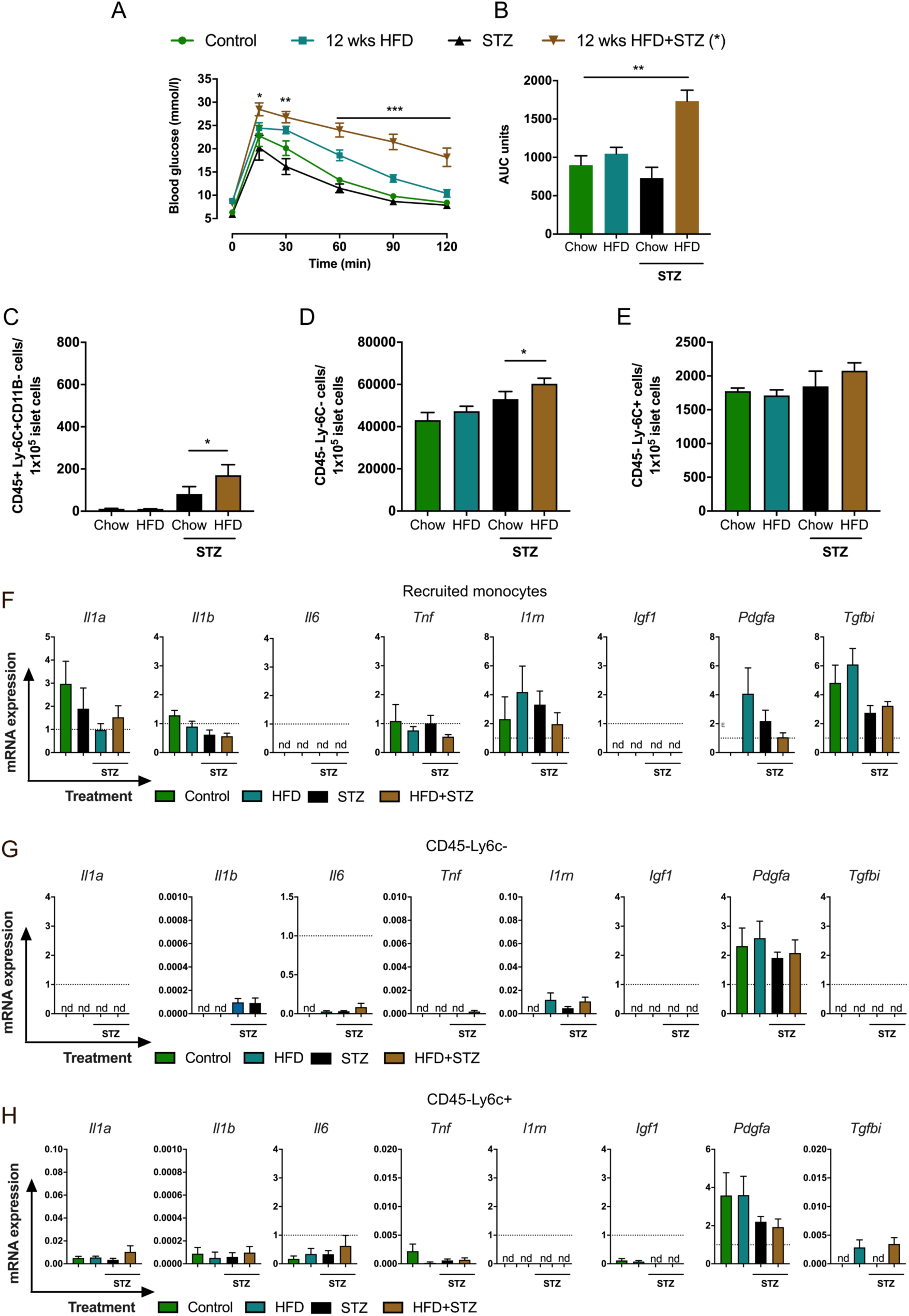
Glucose tolerance, islet immune cell populations and mRNA expression in mice challenged with multiple low-dose STZ + HFD. A) Glucose tolerance test (IPGTT, 1.5 g/kg) one week after administration of the first dose of STZ or vehicle control; n=7-9 mice/ group, *p < 0.05, **p < 0.01, ***p < 0.001 test group versus control, two-way ANOVA with Dunnett’s multiple comparisons test. (B) Incremental area under the curve (AUC) for mice in (A); **p < 0.01 test group versus control, one-way ANOVA with Dunnett’s multiple comparisons test. Fractions of (C) CD45+Ly-6C-cells, (D) CD45-Ly-6C-cells, (E) CD45-Ly-6C+cells from mice described in Figure 5; n=5 for control, STZ groups; n=5-6 for 12 weeks HFD, 12 weeks HFD+STZ groups. * p<0.05, Student’s t-test. (F-H) qPCR of recruited monocytes and non-immune cells. Relative mRNA expression levels of *Il1a*, *Il1b*, *Tnf*, *Il6*, *Il1rn*, *Igf1*, *Pdgfa*, and *Tgfbi* expressed as fold over islet macrophage control; n=4-6. For each sorting sample (n) islets from 3 mice were pooled together (average of 828 +/- 164 islets).

**Supplementary Fig. 7:**
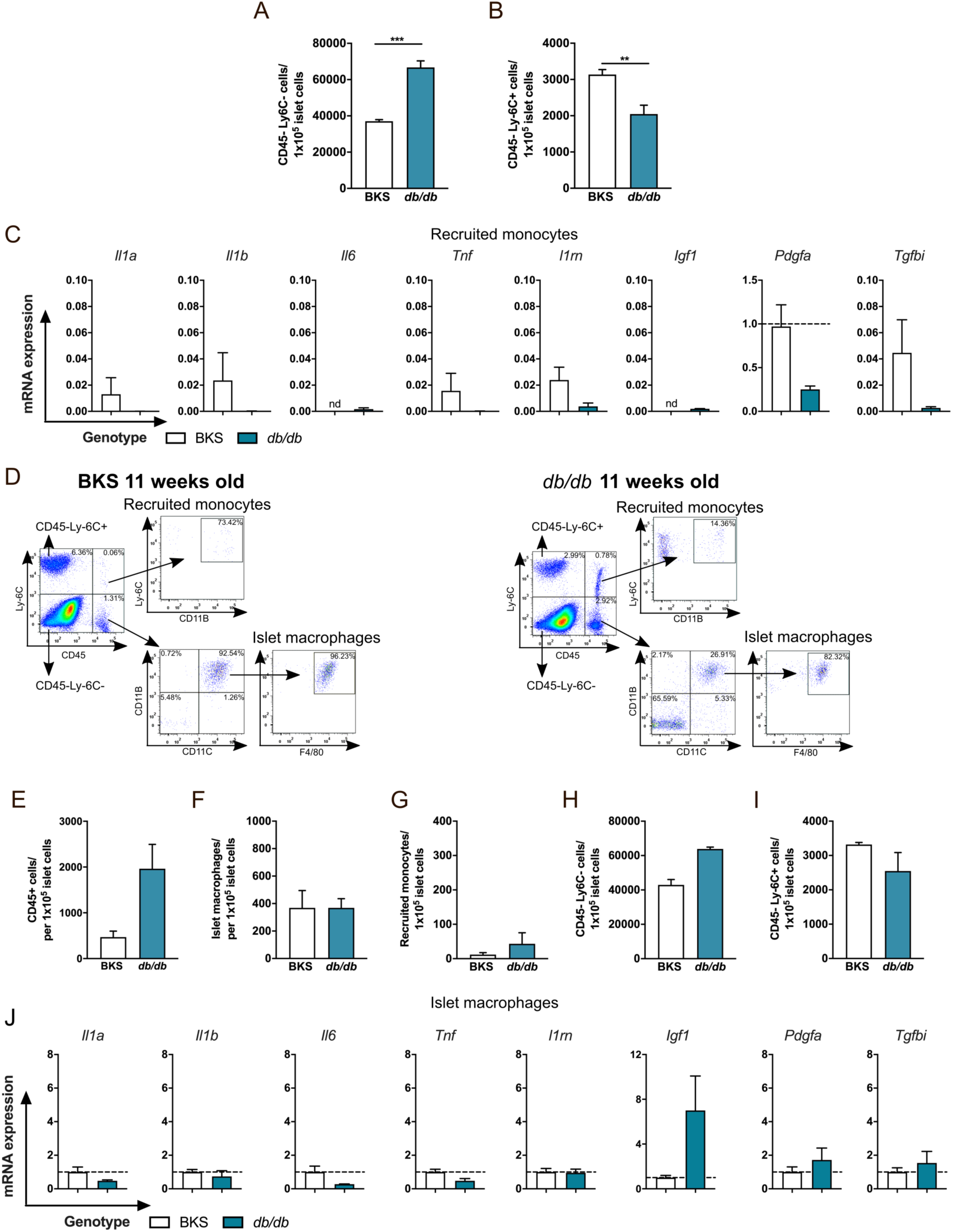
Gene expression and islet immune cell populations in 8- and 11-week old diabetic db/db mice. Fractions of (A) CD45-Ly-6C- cells and (B) CD45-Ly-6C+ cells in islets of 8-week-old BKS and *db/db* mice; (A-B) n=4, 2-4 mice pooled to obtain 556 +/- 52 islets per sample (n); **p < 0.01, ***p < 0.001 *db/db* versus BKS, Student’s t test. (C) qPCR of recruited monocytes from 8-week-old BKS and db/db mice. Relative expression levels of *Il1a*, *Il1b*, *Tnf*, *Il6*, *Il1rn*, *Igf1*, *Pdgfa*, and *Tgfbi* presented as fold control BKS islet macrophages; n=4, 2-4 mice pooled per sample (n). (D) Representative flow cytometry profiles and gating strategy for cell sorting of dispersed islets from 11-week-old BKS and db/db mice. Fractions of (E) CD45+ cells, (F) islet macrophages, (G) recruited monocytes, (H) CD45-Ly-6C-, and (I) CD45-Ly-6C+ cells cells in 11-week-old BKS and db/db mice. (J) qPCR of islet macrophages. Relative expression levels of *Il1a*, *Il1b*, *Tnf*, *Il6*, *Il1rn*, *Igf1*, *Pdgfa*, and *Tgfbi* shown as fold control (BKS). For BKS mice, 2 mice per sample were pooled to obtain 520 +/- 26 islets and 4 sorting samples in total, n=4; for *db/db* mice, 4-5 mice were pooled to obtain 464 +/- 127 islets and 2 sorting samples in total, n=2.

**Supplementary Table 1:**
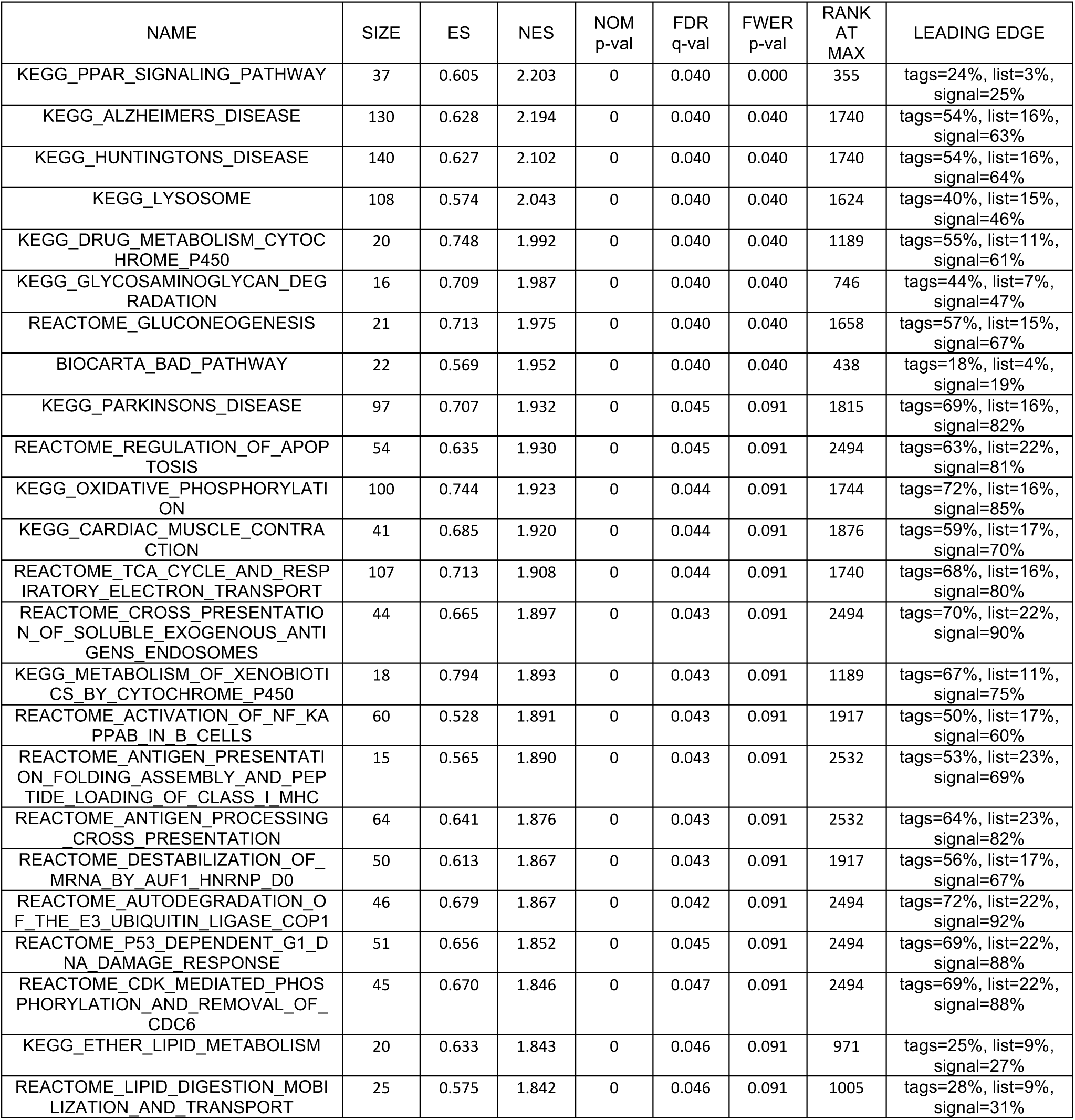
Gene sets enriched in STZ versus control treated islet macrophages with FDR q-value < 0.05.

**Supplementary Table 2.**
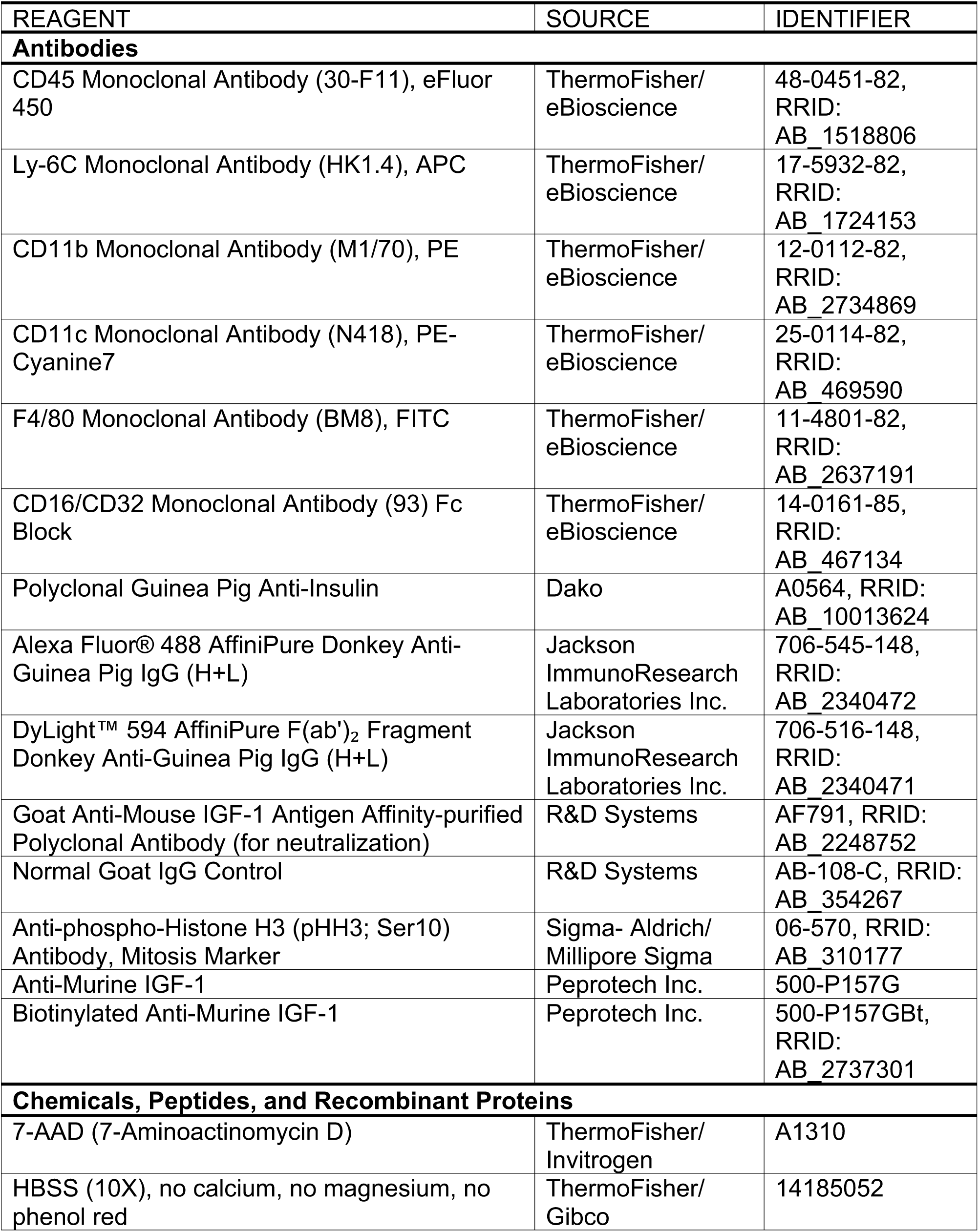
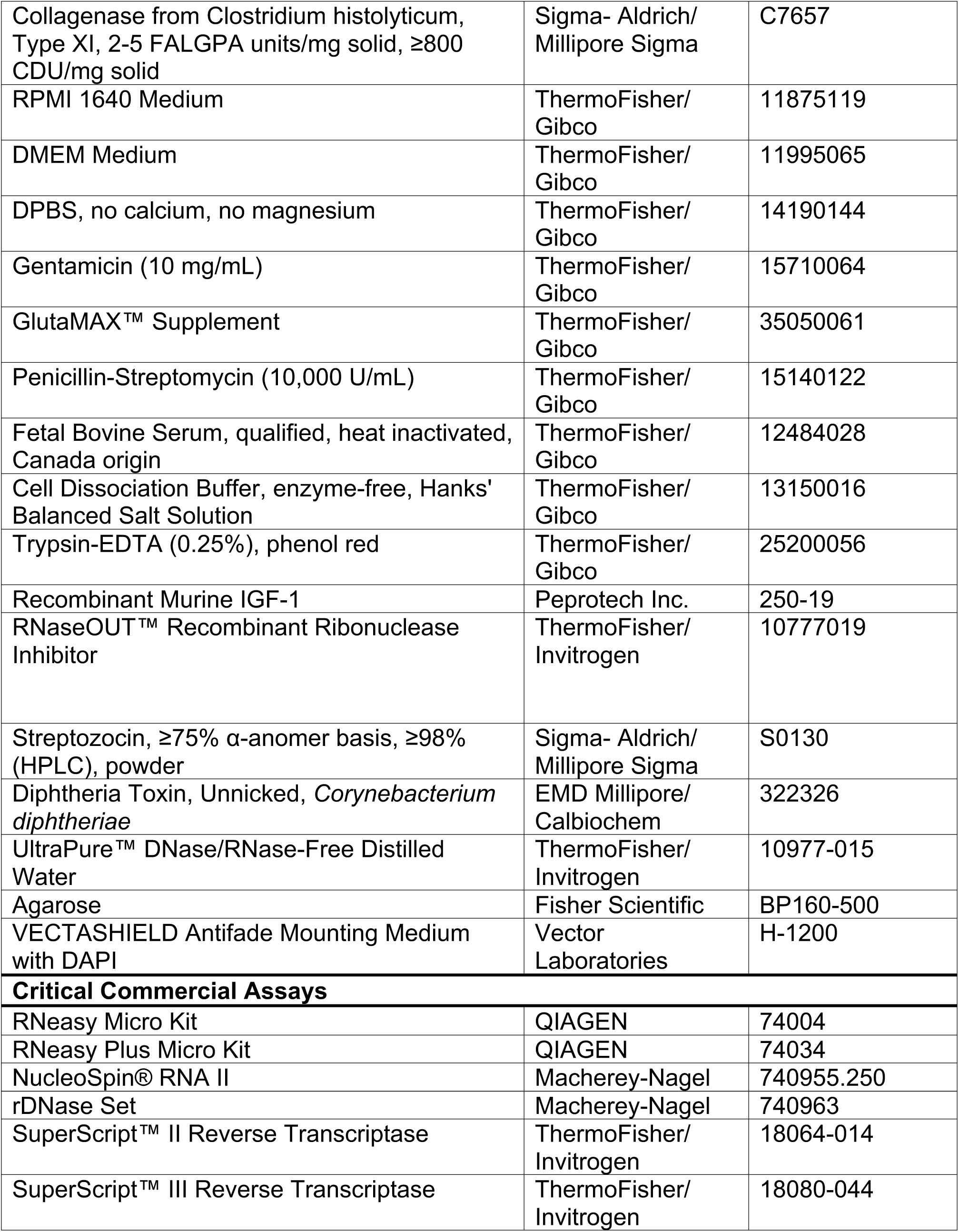
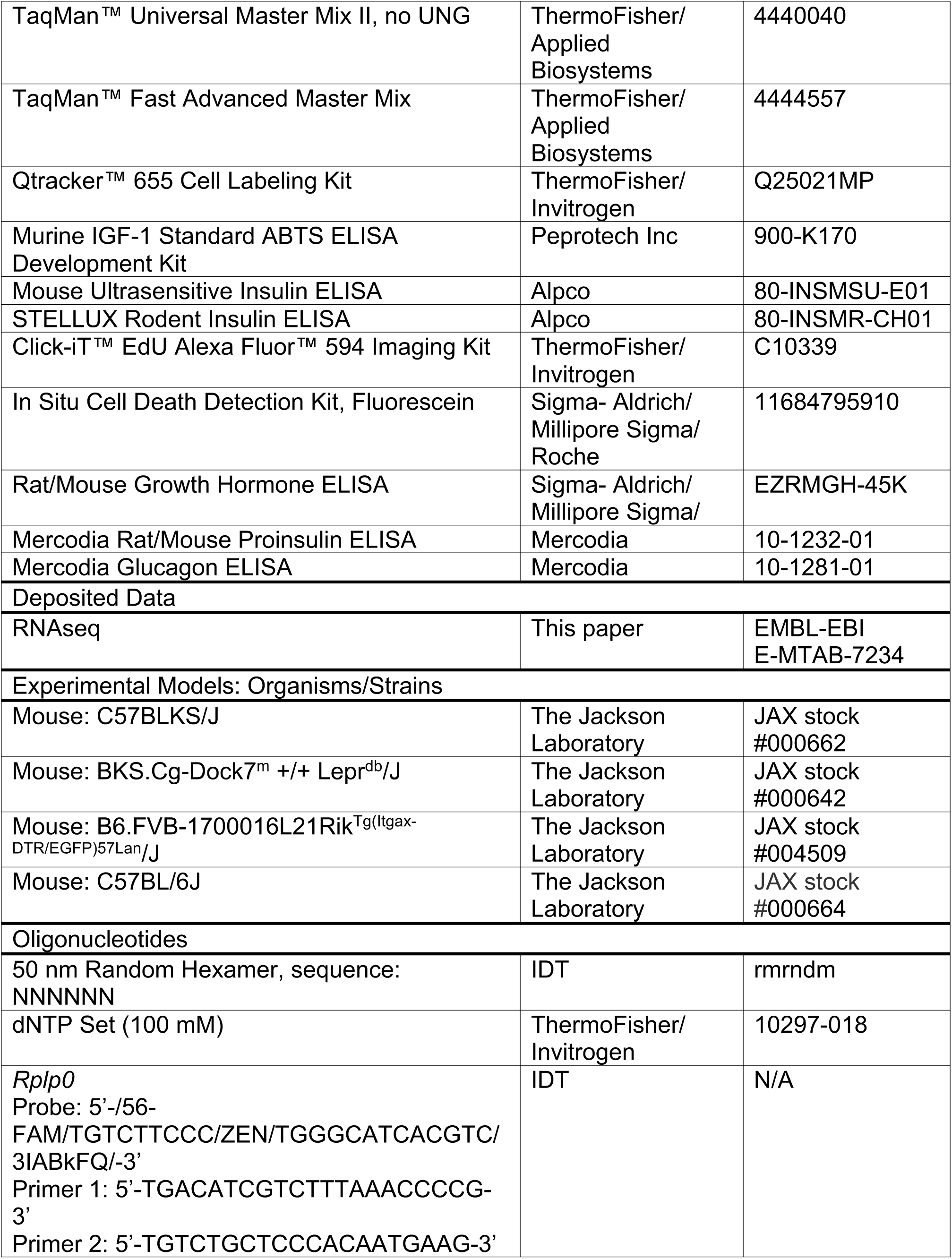
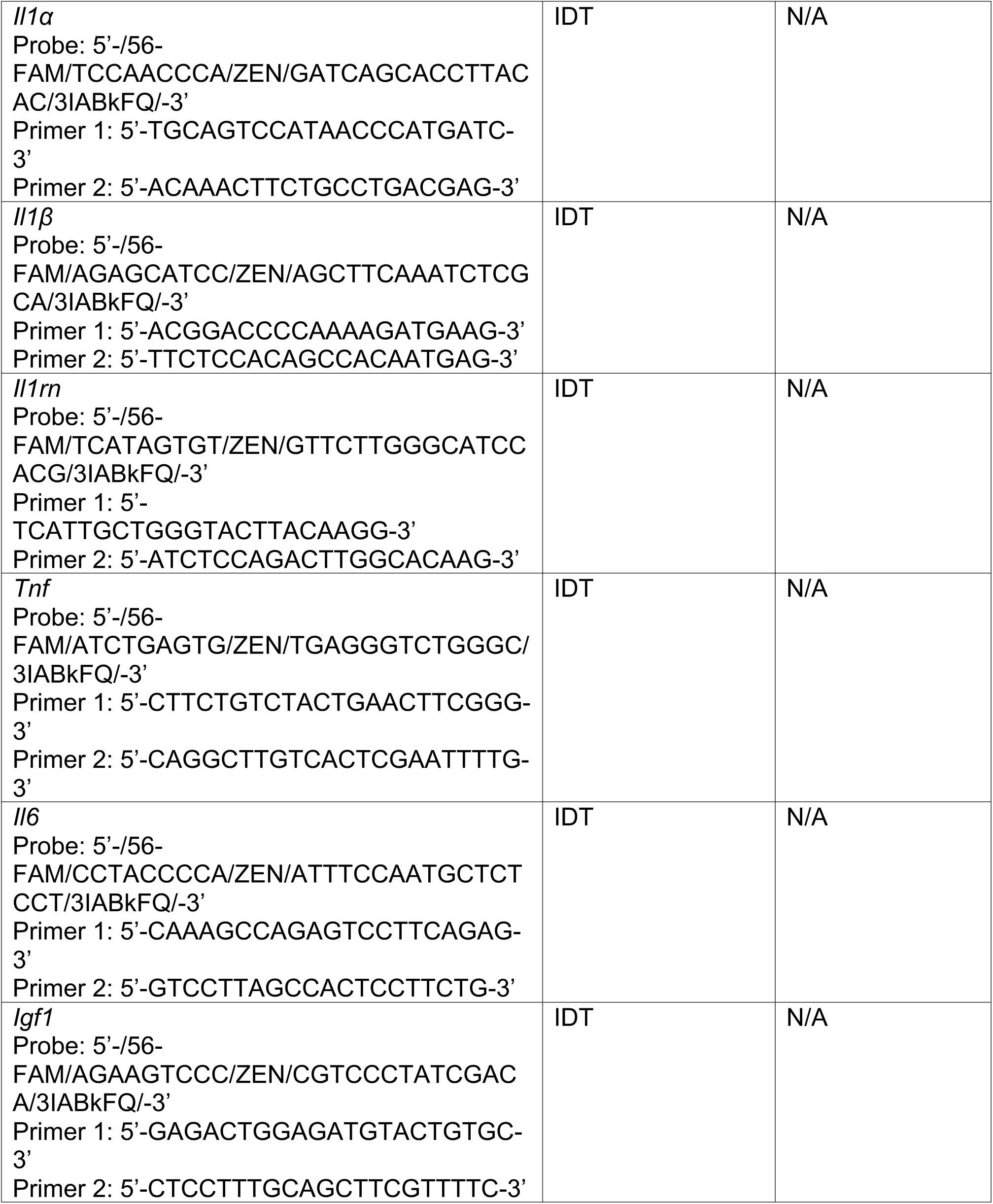
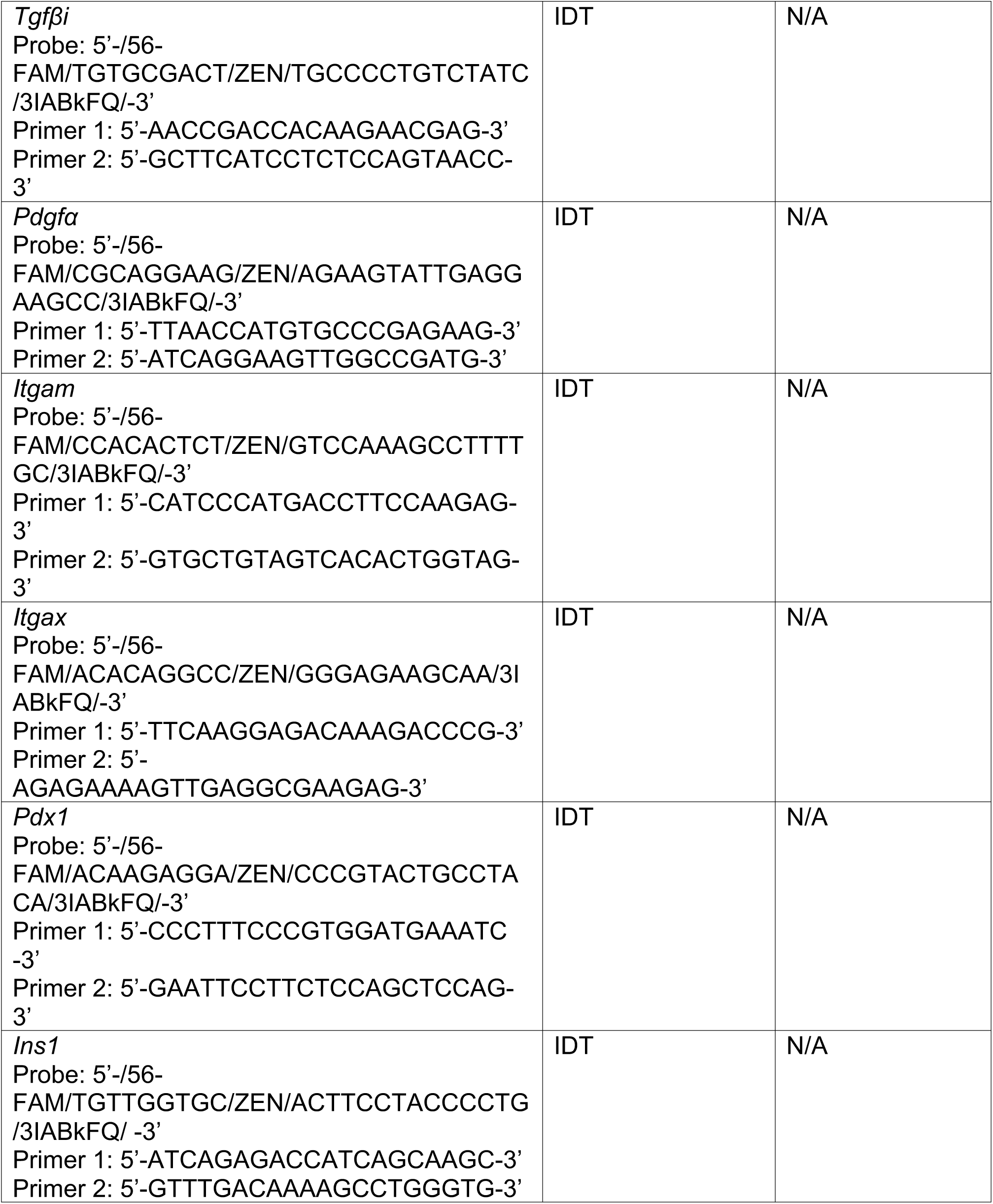
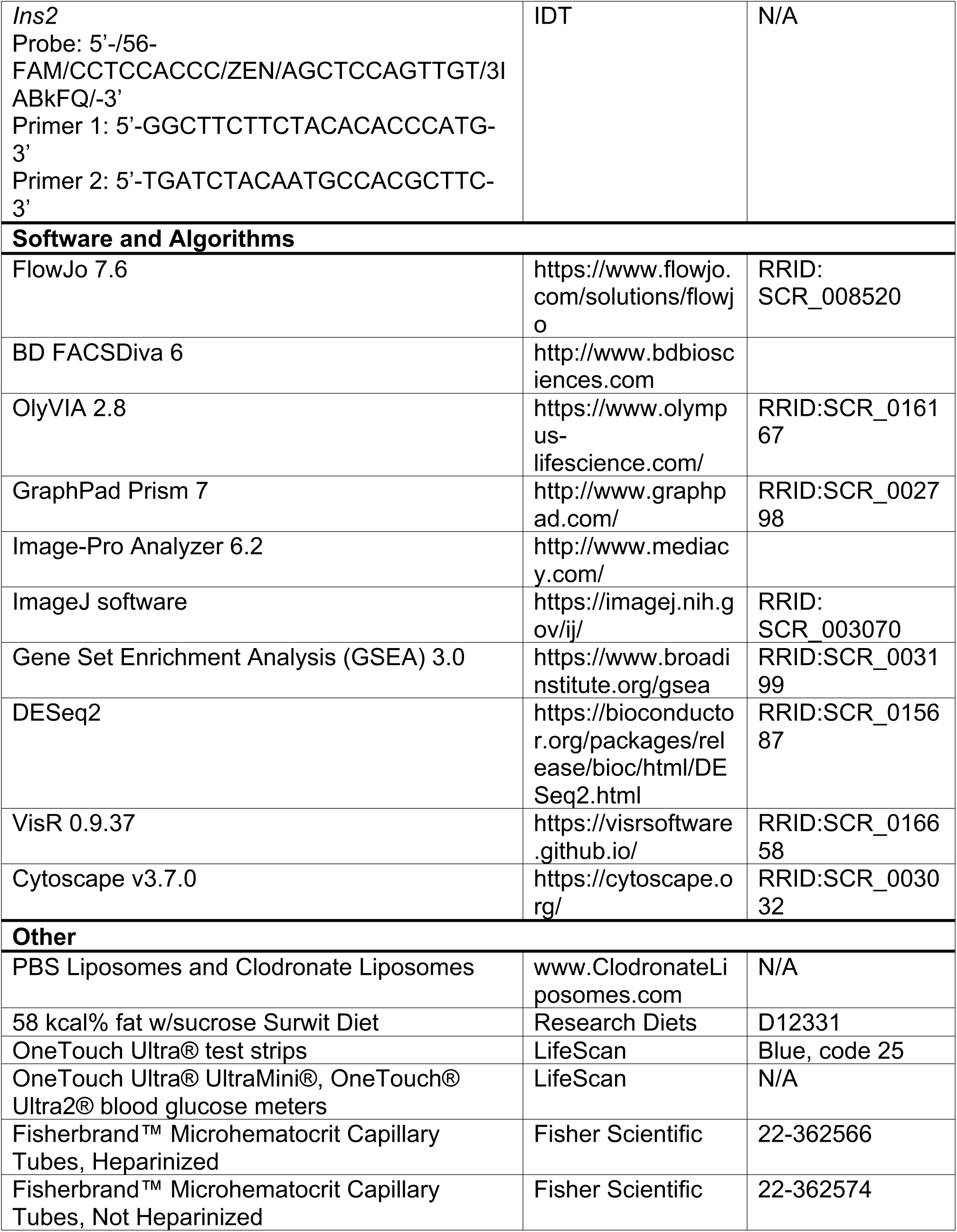
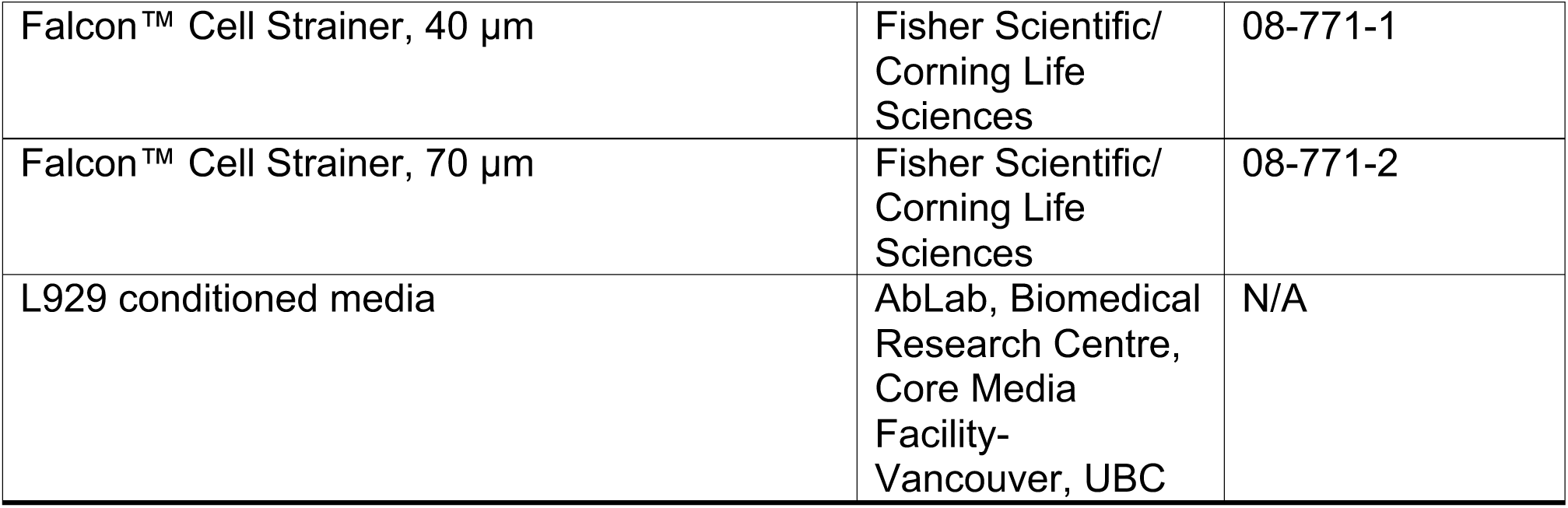
Commercially available reagents and mouse strains used in this study.

